# Is the genomics ‘cart’ before the restoration ecology ‘horse’? Insights from qualitative interviews and trends from the literature

**DOI:** 10.1101/2021.08.05.455206

**Authors:** Jakki J. Mohr, Peter A. Harrison, Jessica Stanhope, Martin F. Breed

**Author notes:** Author for correspondence: Martin F. Breed. College of Science and Engineering, Flinders University, Bedford Park, SA, Australia 5042.

## Abstract

Harnessing new technologies is vital to achieve global imperatives to restore degraded ecosystems. We explored the potential of genomics as one such tool. We aimed to understand barriers hindering the uptake of genomics, and how to overcome them, via exploratory interviews with leading scholars in both restoration and its sister discipline of conservation – a discipline that has successfully leveraged genomics. We also conducted an examination of research trends to explore some insights that emerged from the interviews, including publication trends that have used genomics to address restoration and conservation questions. Our qualitative findings revealed varied perspectives in harnessing genomics. For example, scholars in restoration without genomics experience felt genomics was over-hyped. Scholars with genomics experience emphatically emphasised the need to proceed cautiously in using genomics in restoration. Both genomics-experienced and less-experienced scholars called for case studies to demonstrate the benefits of genomics in restoration. These qualitative data contrasted with our examination of research trends, which revealed 70 restoration genomics studies, particularly studies using environmental DNA as a monitoring tool. We provide a roadmap to facilitate the uptake of genomics into restoration, to help the restoration sector meet the monumental task of restoring huge areas to biodiverse and functional ecosystems.

## Introduction

> “*We’re looking for any tools available that can help us find solutions [to understanding ecosystem processes and functions]. And, genomics provides, you know, a new set of glasses to look at and understand our systems, and therefore, to seek solutions from*” (Scholar 4 in our study).

Humans are now the dominant force in nature, having caused substantial degradation to ecosystems globally, and significant biodiversity loss [1, 2]. The ecosystem restoration sector is tasked with reversing humanity’s ecological footprint by returning and reinstating lost ecosystem services, ecological processes and biodiversity [3], the scale of which is enormous, as highlighted by the United Nations declaration of the *Decade on Ecosystem Restoration* (https://www.decadeonrestoration.org/). The global community have pledged to restore over 350 million hectares of degraded land by 2030 under The Bonn Challenge [4]. Thus, there is great urgency to the upscaling of ecological restoration interventions, from local levels to entire landscapes.

Restoration ecology is experiencing a monumental transformation as it wrestles with challenges posed by climate change, ambitious restoration mandates, and social and political considerations. Tackling such global challenges is no easy feat and requires innovation, science-informed practice, and drawing knowledge from sister disciplines for insight [5–7]. New techniques and methods, such as genomics, offer potential to address restoration challenges [8], but they also invite debate and controversy (see a general description of genomics in the next paragraph, and Box 1 for a technical definition of genomics). Pioneering techniques, like genomics, may be viewed as unproven, and therefore risky by the majority of the field [9], resulting in a significant lag on the uptake of innovation.

### Box 1.

**Definitions and inclusion/exclusion criteria**

*Conservation*: “The protection, care, management and maintenance of ecosystems, habitats, wildlife species and populations, within or outside of their natural environments, in order to safeguard the natural conditions for their long-term permanence” [56].

*Restoration ecology*: “the science that supports the practice of ecological restoration, and from other forms of environmental repair in seeking to assist recovery of native ecosystems and ecosystem integrity” [3].

*Genomics*: Generated *de novo* genomic data using modern sequencing approaches (i.e., high-throughput sequencing; non-Sanger sequencing methods), or that used genomic datasets that had already been generated and ran comparative genomic analyses. Examples of genomics include studies that did (a) *population genomics*, where high-throughput sequencing is used to genotype genome-wide molecular markers (e.g., SNPs) or assemble transcriptomes and/or genomes and make population-scale comparisons; (b) *eDNA analyses*, where high-throughput sequencing is used to characterise particular taxonomic groups (e.g., sequencing 16S rRNA amplicons to characterise bacterial communities) or metagenome-assembled genomes (we included ‘amplicon sequencing’ studies in this category); (c) *comparative genomics*, where the characteristics of entire genomes are compared across individuals within a species and/or across species. See Box 2 for details of population genomic and eDNA examples.

Studies were *eligible* for inclusion where they were:

- published in English language
- published in full text (e.g., not abstracts only)
- published in a peer-reviewed journal [as per UlrichsWebTM Global Serials Directory; 57]
- used genomics for either conservation or restoration ecology, or that propose methods that could be applied to conservation or restoration ecology (i.e., methods with potential implications specific to conservation or restoration ecology, or provided genomic data on a target species)

Studies were *excluded* where they were:

- reviews, perspectives, or essays
- studies of plastid genomes (e.g., chloroplast or mitochondria), or
- studies that used genomics to develop microsatellite markers

*Genomics* can refer to either analysing very large collections of genes within an organism or organisms (i.e., studying genomes or genome-wide molecular markers) or the use of high-throughput DNA sequencing technologies (i.e., genomic sequencing methods, also known as next-generation sequencing methods, as opposed to Sanger sequencing methods). Examples of the two most common applications of genomics used in conservation and restoration include population genomics and environmental DNA (eDNA). *Population genomics*, where very large collections of genes are studied at the population level, can provide a range of very accurate estimates of, for example, the levels of genetic variation, inbreeding, genetic connectivity and demographic histories of these populations. This genomic information can be used to help estimate the capacity of a population to adapt to environmental change. *Environmental DNA* (*eDNA*) methods refer to studying the DNA that organisms leave behind in environmental samples, such as soil or water. The DNA in these samples could, for example, be studied to remotely monitor the presence of invasive or endangered species, or to study entire ecological communities for organisms that are very challenging to count via traditional field-based methods (e.g., microbes).

Disciplines often experience controversy when new approaches challenge existing paradigms and existing practices [10]. Changes in methods can take years to unfold, as information gets disseminated, new practitioners trained, and the conventional wisdom retired. In addition to barriers from status-quo thinking, innovations may require significant financial investments that are often perceived as risky given that the viability of the innovation may not be immediately realised [11]. Indeed, in some situations, new models and paradigms may prove to be less beneficial than originally expected, have unintended consequences, be un-scalable, or be superseded by even better techniques [12]. Further, innovations often require collaborations across disciplinary and geographic boundaries, which can present barriers to progress and delayed uptake [13].

The use of genomics in conservation has faced significant barriers [14], contentious debates on its role [15, 16], and unresolved technical debates (e.g., adaptive vs. neutral diversity [17]; drift vs. local adaptation [18]; species-population continuum [19]); however, despite these challenges, genomics has made a considerable impact on the field of conservation [20–23]. Among the first applications in conservation was population genomics [24], which offered conservation scientists and practitioners detailed insight into species lineages and population demographics. Such data are central to precise delineation of conservation units and key to conservation management [14, 25]. Genomics also has been applied to detect and monitor species of conservation concern [26].

In contrast, restoration is yet to leverage genomics to a similar degree [8]. However, to achieve its ambitious targets, restoration must adopt new techniques and approaches [6, 27]; genomics may be one such technique. Genomics is a valuable and cost-effective technology that, if used appropriately, could greatly advance restoration [8]. To better understand the barriers hindering the uptake of genomics, and possible strategies to overcome them, we aimed, firstly, to understand how leading scholars view the potential role of genomics in restoration ecology. We also explored what insights scholars in restoration ecology’s sister discipline of conservation could offer, a field that has leveraged genomics to address many important issues [14, 28]. Secondly, we explored the trends in publications that have used genomics to address restoration or conservation. Addressing these two objectives, we provide a roadmap to facilitate and leverage genomics to meet global restoration targets.

## Methods

### Qualitative interviews

To better understand the nexus between genomics and restoration, we used qualitative research. Qualitative research methods are appropriate when the phenomenon of interest is complex or poorly understood [29, 30], and such methods are used in both restoration ecology and conservation [31, 32]. We used in-depth, semi-structured interviews to explore scholars’ perceptions of (a) the current and potential role of genomics in restoration and conservation, and (b) perceptions of the barriers to and enablers of using genomics in these disciplines.

We used purposeful sampling, which is a method that identifies study participants based on their ability to provide rich information on the phenomenon of interest. We identified both restoration ecology and conservation scholars based on their background and experience [33]. We sought scholars of different career stages (based on their publication records and scholarly reputations; Table 1), including both scholars who integrate genomics into their studies and those who do not.

**Table 1.** Scholar number, years since PhD, discipline expertise and citation and publication data of respondents (citation and publication data from Google Scholar [accessed 1 July 2020]).

| Scholar | Years since PhD | Conservation or restoration | Genomics used in Research | Citations | Publications |
| --- | --- | --- | --- | --- | --- |
| 1 | >30 | C | Y | >10,001 | >100 |
| 2 | >30 | R | N | >10,001 | >100 |
| 3 | >30 | R | N | >10,001 | >100 |
| 4 | >30 | Both | Y | 5,001-10,000 | >100 |
| 5 | 10-29 years | Both | Y | 5,001-10,000 | 50-100 |
| 6 | 10-29 years | R | N | 1,500-5,000 | 50-100 |
| 7 | 10-29 years | R | Y | 501-1,499 | 50-100 |
| 8 | <10 years | C | Y | 1,500-5,000 | 50-100 |
| 9 | <10 years | Both | Y | <500 | <49 |

Our in-depth interviews included four senior scholars with ≥30 years’ experience, three mid-career scholars with 10-29 years’ experience, and two early-career scholars with <10 years’ experience (Table 1). Respondents were located in North American and Oceania, with their training/background from North America, Oceania and/or Europe. Each respondent was assigned a number for anonymity. Of the respondents, six were genomics experts, while three were familiar with genomics, but did not conduct genomics research themselves. Four respondents worked primarily in restoration ecology, two primarily in conservation, and three conducted research that spanned both conservation and restoration. Our purposive sampling approach is suitable for our exploratory objective but is not meant to be representative of all the combinations of expertise across the two disciplines.

Interviews were conducted between July and September 2019. Prior to the interview, a brief script was read to the respondent, adhering to informed consent guidelines, per Institutional Review Board standards (lead author’s institution #144-19). An interview guide (see Appendix 1 in the Supplementary Information for details) ensured comparability across interviews. We informally tested the interview guide with colleagues and at conferences, prior to submitting it for our ethics board approval. Respondents answered questions about their academic background, their understanding of genomics, how genomics was or might be used in their research field, as well as about their perceptions and experience regarding barriers and enablers. All interviews concluded with a set of reflective questions related to cross-pollination from conservation to restoration ecology, as well as advice to “restoration ecologists interested in increasing the uptake of genomics”. The interviews had a conversational quality; respondents guided the flow of the discussion. Three researchers participated in all interviews, with one of the team serving as the lead interviewer. The other two researchers were invited to interject questions to clarify and probe at selected spots in the interview. This semi-structured approach provided the benefits of organization and flexibility, while minimising the risk of interviewer-induced bias [34]. Each interview lasted about one hour, was recorded, and professionally transcribed verbatim.

Qualitative data were analysed using an iterative process to identify themes emerging from the data [35]. The three research team members involved in the interviews each analysed three-to-four transcripts in detail for insights and themes. We then shared our individual interpretations, discussing discrepant views, and cross-checking themes against the transcripts. Based on this iterative process, we organized the findings into key themes [i.e. ‘thematic analysis’; 36]. Within these themes, we present findings that convey the full range of insights and perspectives within that theme. We then shared our themes and interpretations with the research participants in a written form to ensure the findings resonated with their experience and perceptions. The themes that survived this validation process are discussed in the results below. Data excerpts illustrate specific findings and provide empirical evidence for the themes that reflect our research aims. During our interviews, we asked additional questions, such as the respondents’ perceptions about the relationship between conservation and restoration. These additional questions ensured respondents felt insights from one discipline to another were relevant and provided context for our interpretation of their responses; however, we do not report these additional findings here.

### Examination of research trends

We investigated trends in published research regarding the use of genomics in restoration ecology or conservation (see Box 1 for definitions). The question posed was *what are the temporal and geographic trends in the published studies using genomics in conservation and/or restoration ecology studies*? We searched two library databases on March 1, 2020 to identify all peer-reviewed articles published in English language that used genomics within conservation and/or restoration ecology studies. The search was performed in Web of Science Core Collection and Scopus (see Appendix 2 for details).

Results from the database searches were exported into Endnote X9 for management. Within Endnote, duplicates were manually removed, then the titles and abstracts were screened by two authors for inclusion/exclusion, based on predetermined criteria (Box 1). The full texts of all remaining articles were obtained and screened against the same criteria. Where there were uncertainties regarding inclusion of a study, another author independently assessed the study.

The following data were manually extracted from all included articles into a Microsoft Excel spreadsheet: country of the study, year(s) of data collection, publication year, whether the study related to conservation and/ or restoration ecology (based on the definitions in Box 1), and the genomic methods used (e.g., eDNA, population genomics). A second author independently coded the data from any study where the other author was uncertain. Characteristics of the included studies were reported descriptively and graphically [37].

## Results and Discussion

Our qualitative findings identified three inter-related themes in terms of the barriers to and enablers of using genomics in restoration – academic training and background, methodological considerations, and philosophical perspectives – which we discuss in turn below. We also found common ground among the interviewees: the need for more use cases of genomics in restoration and the value of multi-disciplinary collaboration. The insights gained from these qualitative data prompted us to explore the trends in the database search to better understand where and how genomics tools were being applied in conservation and restoration, which are discussed following the qualitative findings.

### Qualitative Findings

#### Academic training and background

Scholars expressed the importance of background training in genetics and evolutionary biology to harness the power of genomics in restoration. Scholars less experienced in genomics acknowledged this need and expressed concerns that without some training in genetics or evolutionary biology, the stereotype of restoration “*as a gardening exercise*” (Scholar 3) would present a barrier to new perspectives. Scholar 2 (also a non-genomics expert in restoration ecology) stated that the understanding of genetics “*and all that stuff about small populations and genetic bottlenecks is well-embedded in conservation thinking and much less so in restoration*.” Scholar 8, a genomics expert in conservation, noted “*the restoration field is more ecologist-driven than geneticist driven*.” The need for at least some training in genetics or evolutionary biology to harness genomics has been a topic of study in conservation [39]. Similarly, non-genomics experts in our interviews expressed concern about how “*anyone who manages an ecosystem could do so effectively*” (Scholar 6) without training in genetics or evolutionary biology.

Relatedly, genomics experts also acknowledged the skills needed to handle the ‘big data’ aspects of high-throughput sequencing and requisite bioinformatics, describing themselves as “*total data nerds* [who] *dive into two terabytes of data with great enthusiasm*” (Scholar 9). Scholar 9 continued, noting that traditional restoration ecologists may be “*unprepared for the fact that they can’t open this data in Excel and you can’t eyeball it*…*It requires specialist expertise to be able to handle this sort of data*.”

#### Methodological considerations

Our interviews revealed the importance of rigorous designs for interpreting the results from genomics data, which was emphasised by all scholars. For example, Scholar 3 (a non-genomics restoration scholar) suggested that although genomics allows researchers to describe important genetic differences between plant populations faster, “*we still need to ascribe functional significance to those differences*”, stressing the need for classic experiments with control groups in order to “*close the loop on the functional impacts of genomics findings*.” Genomics expert, Scholar 7, shared this view, noting that: “*People who embrace new technologies can forget the old school methods of just growing plants in a glasshouse*.” Similarly, genomics expert, Scholar 5, noted that with respect to genomics studies, “*we’re still analysing and trying to figure out the best analyses to take for these big* … *genomics studies. The challenge is, even with the expertise, getting the analysis right, making decisions around which comparisons and what analysis do you run*.”

In addition, scholars expressed their thoughts about the confidence in drawing conclusions based on methods developed using model organisms. The field of conservation benefited from publicly-available genomic resources, such as assembled and annotated genomes stemming from human medicine or agriculture [40]. “*Most of the first examples of using genomics in conservation benefitted from agriculture, where they had already collected and analysed the genomic information*” (Scholar 1). Experienced genomics scholars were comfortable using these model organisms. Scholar 5 stated, “*As an evolutionary biologist, I can say fundamentally, how genetic diversity drives population dynamics and persistence is universally the same, regardless of taxa*.” These more experienced genomics scholars wondered whether the less experienced genomics scholars “*are reticent to accept arguments about the importance of considering genetic diversity when that data comes from model organisms. [They wonder how] does that translate to my particular species or my ecological context?” (Scholar 5)*.

Another methodological consideration was whether genomics data could be used to explain genetic variation between populations that is relevant to restoration. Experienced genomics scholars were comfortable using genomics to understand the genetic basis of adaptation. For example, Scholar 9 said, “*As much as anything, I’ve been following my nose; genomics can help us take more apart and start to understand adaptive differences*”. Still, restoration ecologists with less experience in genomics expressed concern about the inability to draw precise conclusions from genomics data. Scholar 3 noted that “*genomics is important but it’s not everything*,” and wondered whether genomics researchers might be putting the “*cart before the horse*.” Wondering about whether the cart might be before the horse leads to the third theme in our qualitative findings.

#### Philosophical perspectives

Another area of findings, related to the prior two, is the scholars’ attitudes and beliefs about embracing both the upside potential and downside risks that come with using genomics in restoration. We refer to this as their philosophical perspectives about innovation. More classically-trained restoration ecologists expressed concern about genomics scholars ‘overpromising’ what genomics can do for the field, that genomics was ‘over-hyped,’ genomics scientists “*have the hammer and are looking for the nails*” (Scholar 3). Some scholars with less experience in genomics characterised other restoration ecologists as being risk averse: “[some] *restoration people don’t like anything new at all*…” (Scholar 2), as well as their possible concerns about how “*scary and troublesome*” genome editing is: “*if you unleash this technology into nature, you’re unleashing the ‘dogs of hell’ and it’s all going to be bad*” (Scholar 2). This finding is consistent with innovation theory [41], which highlights that later categories of adopters are more cautious and sceptical about novel innovations; they worry that novel innovations may be unsafe or have unintended consequences.

Scholars experienced in genomics expressed caution on the use and interpretation of genomics data. For example, Scholar 4 wanted to avoid “*putting the cart before the horse*,” with explicit recognition of the work yet to be done to validate genomics’ potential – even if it does not pan out as expected. Despite the need for “*creative thinkers, people willing to take the chance, to jump upon those new methods, and think outside the box*” (Scholar 8), experienced genomics scholars simultaneously acknowledge the need to temper the hype. Scholar 7 said, “*There’s a lot of scepticism on the method* [genomics] *but there’s also a lot of excitement* – *which we’re trying to temper by saying we need to do a lot of research to actually show how well this is going to work*.” Scholar 7 continued, “*People are running to it because it’s new and shiny… we need to set realistic expectations about how well genomics is going to work, how much work we need to put in to prove how well it’s going to work*.” Scholar 8 agreed with the need for tempering: “*I understood how difficult genomics was, how uncertain it was, and I was like, ‘Hold on. Pump the brakes. I’ve worked really hard to get through to the more senior people the difficulty and the uncertainty surrounding these data…It’s not a silver bullet. There’s so much we need to do to make it a usable and reliable, trustworthy tool for restoration assessment*.”

#### Solutions

A final set of qualitative findings regards strategies to address challenges in leveraging genomics for restoration ecology. Firstly, both genomics experts and those less experienced with genomics stressed the need to “*build up enough case studies to demonstrate positive outcomes in terms of the success of particular restoration programs*” (Scholar 5). This scholar emphasised the importance of identifying and validating “*current use cases where genomics could offer tangible value in research today and addressing pressing questions now*.” Similarly, Scholar 3 noted that “*the greatest challenge for genomics is to demonstrate ‘runs on the board*”, to show that it is a cost-effective tool that provides relevant answers to restoration questions that, without which, practitioners may make less optimal management decisions. Our results are consistent with discussions on understanding the attitudes of scientists on genetics and evolutionary principles, which has previously identified the lack of case studies as issues for these topics [42, 43]. However, we note that this stated desire for case studies contrasts with the findings from our examination of research trends (see below), which counted 70 studies that leverage genomics to address restoration issues – evidence of the nascent body of research in using genomics in restoration ecology. A barrier in some restoration contexts in developing genomics case studies is they can take a very long time to validate (e.g., it can take 35+ years to comprehensively analyse local vs. non-local seed sources informed by genomics for long-lived plants).

Related to the need for case studies, the interviewed scholars emphasised the need for greater education and training regarding what genomics is, the types of research questions it can usefully address today, and generally, building capacity, skills and knowledge. Moreover, without improved training in evolutionary biology and genetics, restoration ecologists may find it difficult to harness genomics to address critical questions in climate change adaptation, ecosystem resilience and soil health, for example.

A second recommended strategy scholars emphasised to overcome challenges is the importance of collaboration: between conservation biologists and restoration ecologists, between classically-trained restoration ecologists and genomics experts, and between practitioners and scientists. Scholars 3, 4, and 6 each emphasised the benefits of interdisciplinary approaches and collaborative teams comprising people with very distinct skill sets. Scholars 5 and 7 noted that collaborations between practitioner and scientists are as important as collaborations between genomics experts and classically-trained restoration ecologists. Scholar 5 believed that having joint discussions with practitioners regarding the design of field-based studies could advance uptake and demonstrate the benefits genomics might offer in certain areas of practice (e.g., assessing soil function; for ecological monitoring) to close the gap between science and practice. However, Scholar 3 worried that scientists see the value in the tool, but practitioners will not. This scholar described “*the genomics people*” as “*the guys in the other room*”, who “*have to demonstrate that they understand what the restoration community does*” to bring the conversation about genomics in restoration “*into the fold*.” This us-them distinction could stymie collaborative efforts. Scholar 7, a genomics expert, stated, “*Genomics complements traditional approaches. We’ve got to work together with people who use traditional approaches, work side-by-side with them. Our work will feed into theirs and their work will feed into ours. Together, we will make it better and, you know, have better restoration outcomes for the world*.” This perspective of involving multiple groups of people with different perspectives in the coproduction of knowledge offers the opportunity to build pathways towards addressing ecological, political, and technical challenges [44] – an apt description of both genomics and restoration. Coproduction of knowledge could also help to ensure that genomics does not widen the gap in adoption by practitioners, an issue raised previously in discussions of the value and uptake of genomics in conservation [14]. Hence, we advocate for proactive thinking about the value of iterative processes involving diverse types of expertise, knowledge and actors to produce context-specific knowledge [44] that practitioners find valuable as well.

### Examination of research trends

Our database search identified 1845 unique articles, 176 of which met our inclusion criteria and were included in our review (see Appendix 3 for the flow chart of study inclusion/exclusion; full details of all included papers are in Appendix 4). Based on the field definitions we used in our review (Box 1), 106 articles were classified as conservation only, 35 were restoration only, and another 35 had results applicable to both conservation and restoration. The studies that span both conservation and restoration highlight the potential for bridging the discourses between the two disciplines as identified above (see “Academic training and background”). Indeed, collaborations between the two disciplines can help diffuse the use of genomics and knowledge more broadly.

The finding of 35 studies that use genomics in restoration (in addition to the 35 that span both disciplines) contrasts with the qualitative data that emphasised the need for “*runs on the board*” and “*case studies of restoration using genomics*”. Distilling the reasons for this discord is complex; perhaps the studies employing genomics may not yet represent a viable approach for the restoration discipline. Alternatively, it is possible that ecology scholars focus their literature reading within their own area of expertise and are not exposed to genomics research [45]. Or perhaps the lexicon and presumed knowledge that complicate new technologies may hinder the recognition of value that genomics may offer. Regardless of the reasons, the need for relevant and understandable case studies was voiced by both less and more experienced genomics scholars. Box 2 provides broadly-applicable examples that illustrate the use of genomics in a restoration context. The cross-pollination of specialist journals in these disciplines offer a further potential to increase the visibility of genomics in restoration. We recognise that classically-trained restoration ecologists may find the genomics terminology difficult to understand, and hence, reinforces our suggestions for education and training, and general cross-fertilisation to overcome such lexicon and methodological barriers.

#### Box 2.

**Examples of studies that used genomics to address restoration questions.**

We identified 70 studies from our database search that used genomic methods in a restoration context. Here we provide details of four examples of these studies, where these studies include: (A) the earliest eDNA study (and restoration genomics study in general); (B) the equal earliest population genomics study; (C) a more recent eDNA study that used more advanced molecular methods than typically employed; (D) the only study that combined both eDNA and population genomics.

Ficetola *et al.* (A) [46] used eDNA methods in a laboratory environment to demonstrate how this approach could be used to detect the presence of an invasive frog species in freshwater environments. These findings are important for restoration as detecting aquatic vertebrate species – whether invasive, rare or common natives – is often an expensive exercise that is challenging in certain hard-to-access environments and especially when the organisms are in low abundance.

Steane *et al.* (B) [58] used population genomics to identify adaptations across the range of a tree species that is commonly used in restoration plantings. The use of population genomics here helps to inform seed sourcing decisions that take into account the adaptive variation among populations, which for long-lived plants such as most trees would otherwise require many years of commongarden field trials.

Guo *et al.* (C) [59] used eDNA methods in a field environment to determine how the taxonomic and functional gene diversity and composition of soil microbes had changed after restoration plantings. The monitoring of soil microbial communities in restoration is important as they provide key ecosystem services (e.g. nutrient cycling) and are a rich source of biodiversity in their own right. However, microbial communities are impossible to monitor accurately without the assistance of molecular methods since most taxa are not culturable or easily identifiable. The authors combined a more advanced molecular technique – shotgun metagenomics, where essentially all the genes from all the organisms present in a given sample are sequenced on a high-throughput sequencer – with the more commonly-used amplicon sequencing, where a specific gene is amplified in a given sample and sequenced on a high-throughput sequencer and variation in this gene provides a diversity profile of a particular taxonomic group (e.g., bacteria via 16s rRNA sequencing).

Dittberner *et al.* (D) [60] used a combination of eDNA and population genomics to monitor species hybridisation and population admixture between populations of two Arabis plant species. eDNA was used to identify the two species, and population genomics was used to determine population admixture. Sometimes plant species are challenging to identify using traditional morphological approaches and eDNA approach used here can assist in this process. Measuring gene flow and admixture between populations is extremely challenging without the use of molecular methods, and population genomic methods provide great insight into these aspects of habitat connectivity and adaptive potential of populations.

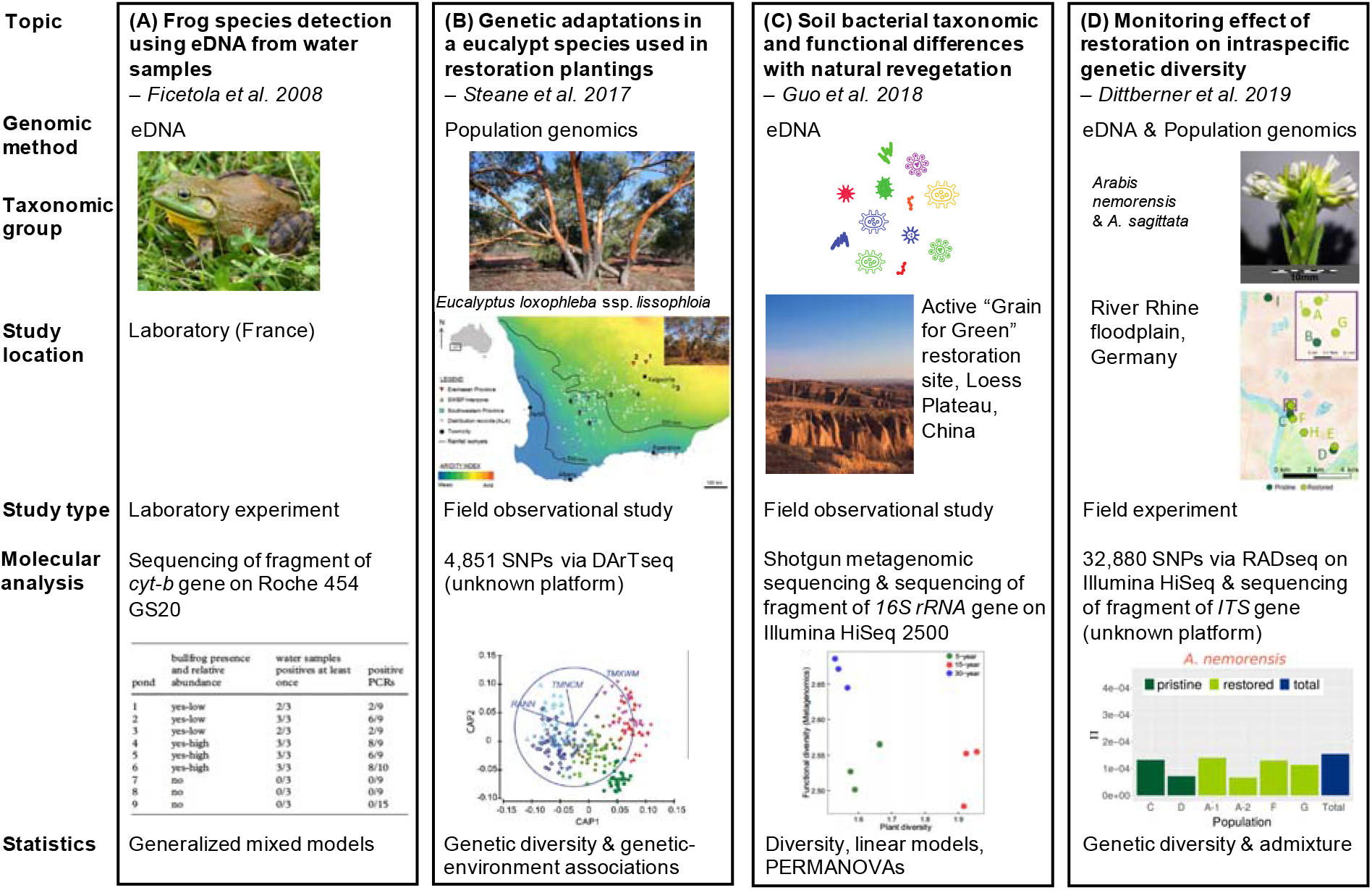

The earliest application of genomics in conservation or restoration detected in our dataset was 2008 [46], followed by a period of very few studies between 2009 to 2012, after which there was an exponential increase in studies applying genomics in both restoration and conservation (Figure 1). Most studies were published between 2018-2020 (59 conservation, 26 restoration, 21 conservation and restoration). This temporal trend appears consistent with other review studies on conservation genomics [47]. The recency of these studies may also help explain the perceived lack of “*runs on the board*” (Scholar 3) as they may not have had sufficient time to become well known. While we counted more studies that used genomics to address conservation issues than restoration, the temporal trend showed a rapid increase in restoration studies that used genomics over the past two years. The temporal trend in growth of publications suggests genomics studies applied in a restoration context may surpass the prevalence of genomics in conservation studies. While the use of genomics in restoration is increasing, this trend might also reflect increased cross-over between the two disciplines.

**Figure 1.**
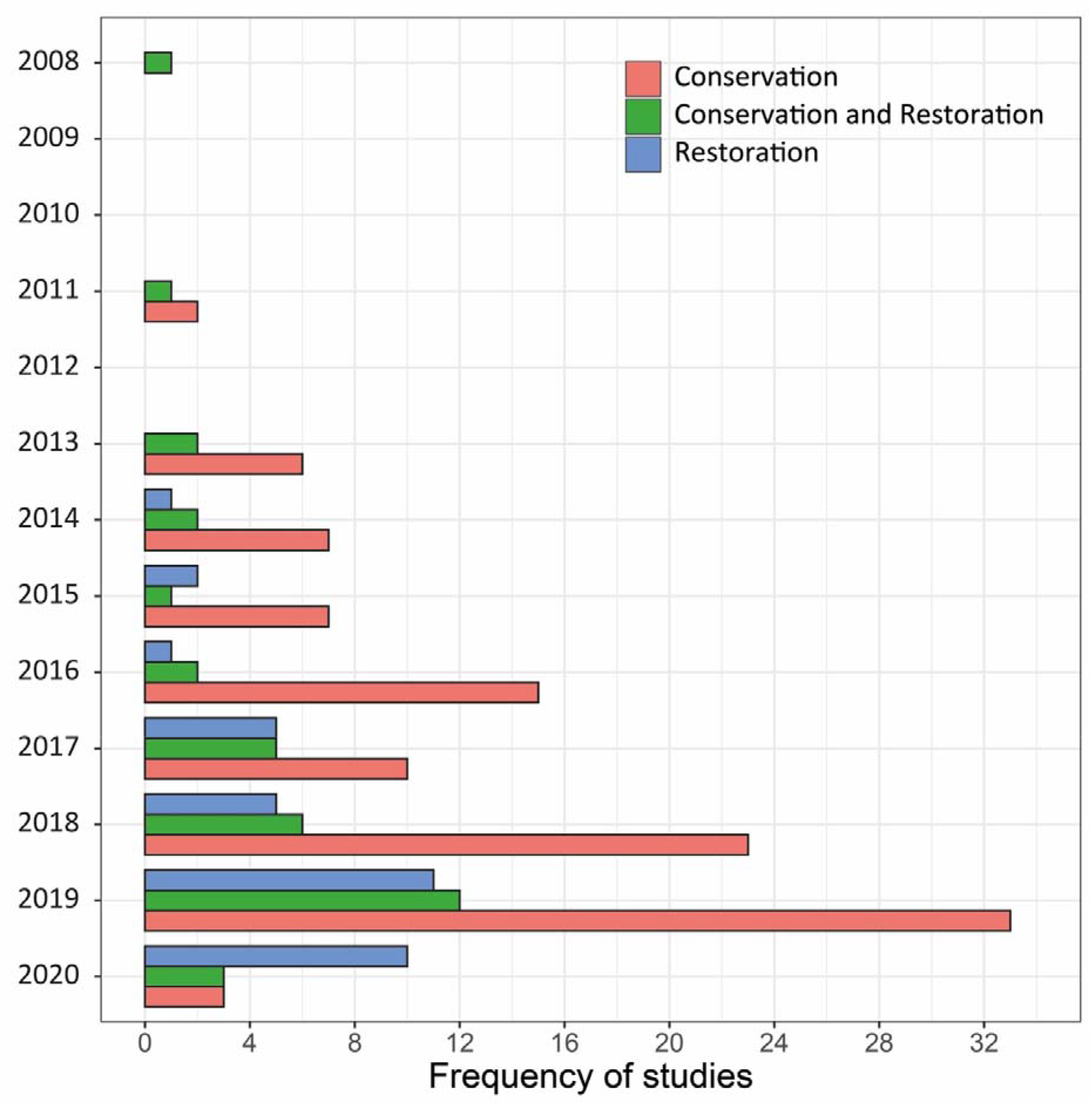
Frequency of published studies over time, from Jan 2008 to March 1, 2020. Studies were divided into conservation (orange), conservation/restoration (green), and restoration (blue).

With respect to geography, most conservation genomics studies were undertaken in North America (n = 36) and Europe (n = 17; Figure 2). These same two continents were among the least common locations reporting restoration genomics studies (Europe: n = 4; North America: n = 8; Figure 2). The number of conservation papers from North America and Europe has generally increased annually, which has not occurred in restoration genomics papers (Appendix 5). However, the number of restoration genomics papers from Russia, China, South Asia shows a clear increase, with the number of restoration studies overtaking the number of conservation studies in 2020 in these locations (Appendix 5).

**Figure 2.**
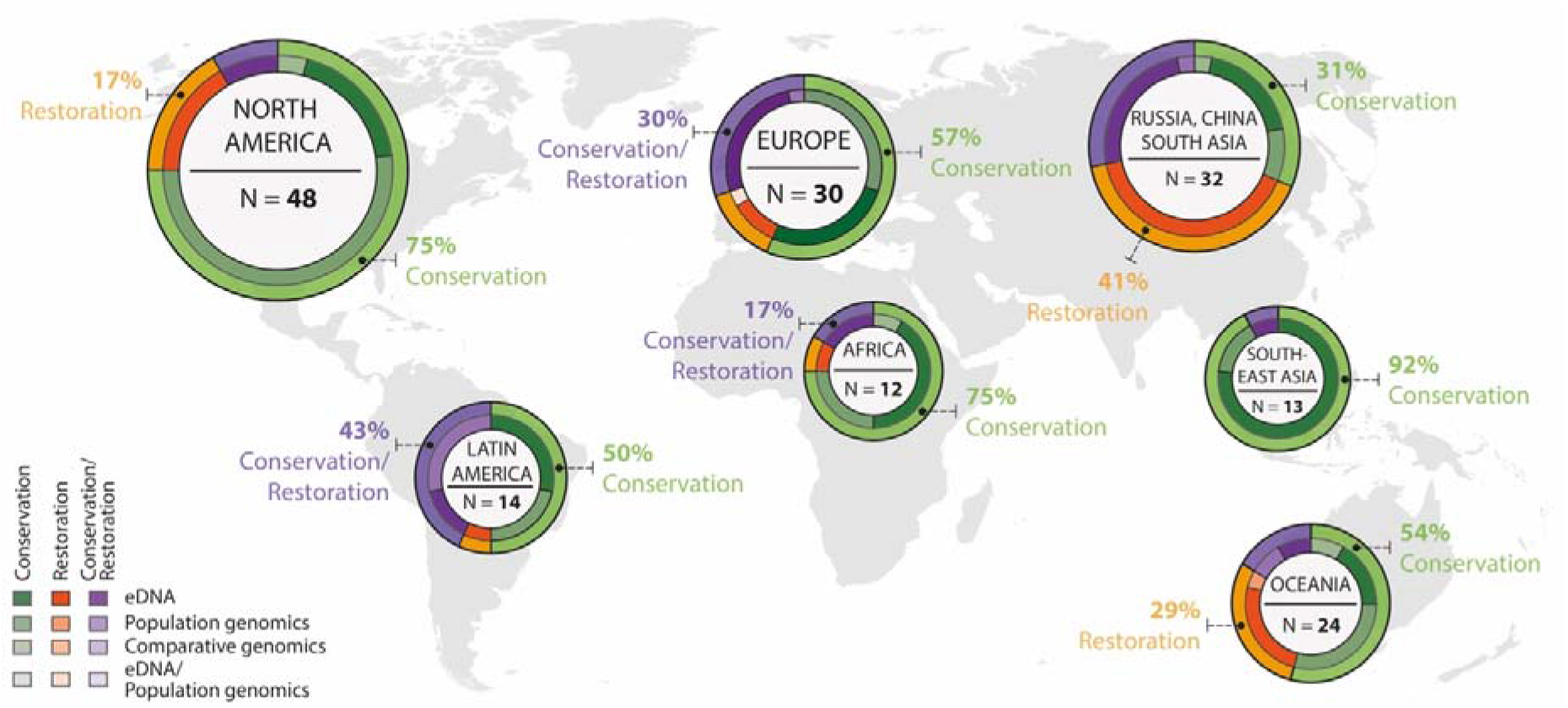
Proportion of studies for seven geographic regions. The outer circle of the donut plot corresponds to the total proportion of studies belong to conservation (green), restoration (orange), and conservation/restoration (purple). The inner circle of the donut plot shows the break-down of each study into the applied genomics method. Studies were assigned to the seven regions based on the country the study was located.

Across the studies assigned to conservation, restoration, and conservation and restoration, the majority of articles reported the use of eDNA approaches (conservation: n = 47; restoration: n = 33, conservation and restoration: n = 27), and a steady increase over time (Figure 3). This genomics approach was most used in conservation and restoration studies from Russia, China, South Asia (n = 27), North America (n = 21), and Europe (n = 20; Figure 2). These findings indicate that there may be broader opportunities in restoration genomics through the appropriate utilisation of eDNA compared to population genomics, as suggested in previous studies [48], but there remains a clear use of population genomics in restoration to inform seed sourcing practices [8].

**Figure 3.**
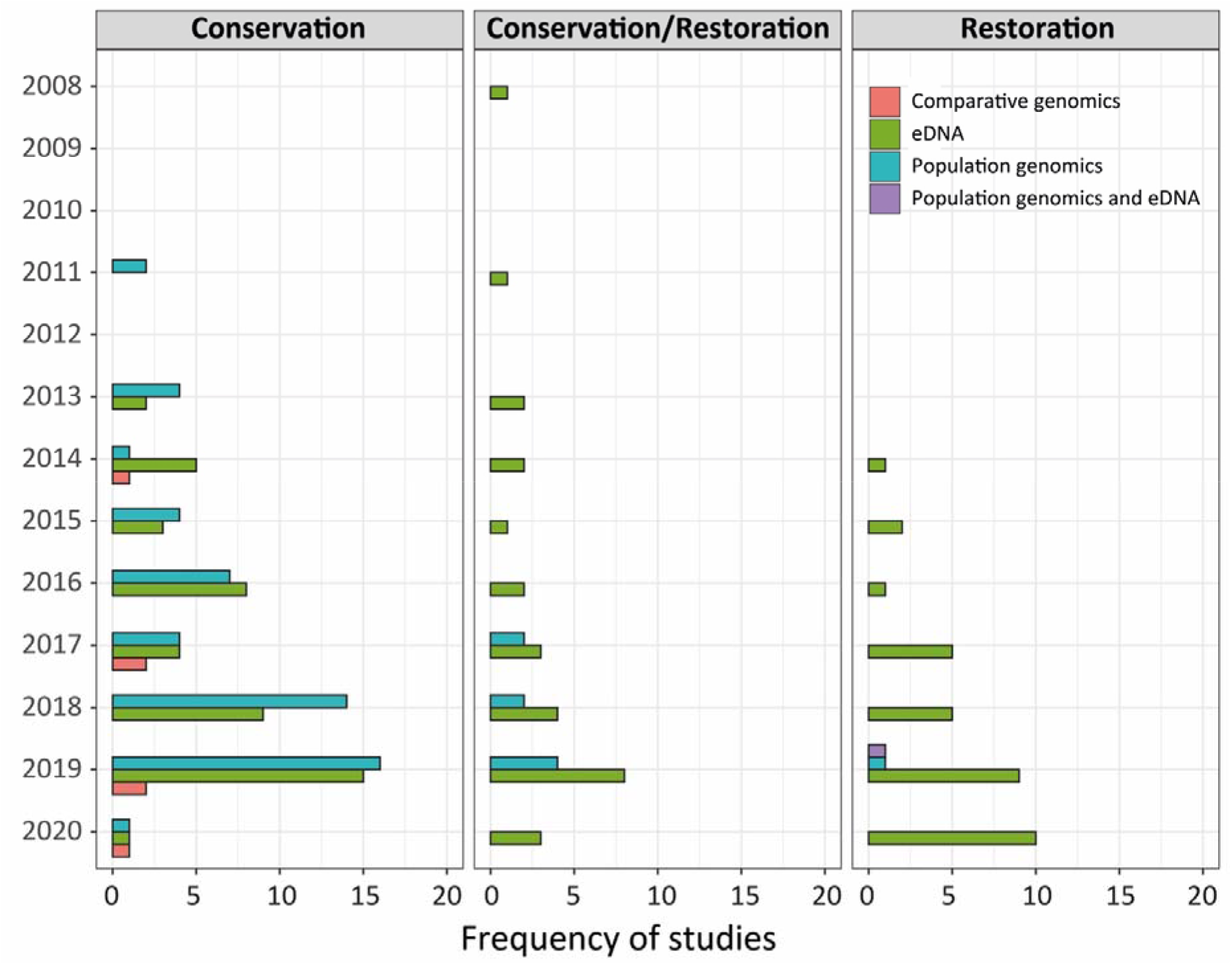
Frequency of published studies across time for conservation, conservation/restoration, and restoration. Studies within each of these divisions were broken down to the applied genomics method, including comparative genomics (orange), eDNA (green), population genomics (blue), and population genomics combined with eDNA (purple).

Population genomics was utilised in 53 conservation studies, 1 restoration study, and 8 studies that crossed over between conservation and restoration (Figure 3). Between 2018-2020, there was a marked increase in the uptake of population genomics, which was most pronounced in the conservation studies (n = 31) and conservation and restoration studies (n = 6; Figure 3). The use of population genomics in conservation was most evident in studies from North America (n = 25); however, we detected no restoration studies from North America that utilised population genomics (Figure 2). Rather, Latin America (n = 4; pooled across restoration and conservation and restoration) and Oceania (n = 3) were the most common locations to use population genomics in a restoration context.

### Helping genomics ‘cross the chasm’ into restoration

A prominent model used to develop a strategy to hasten the market uptake of novel innovations such as genomics is ‘crossing the chasm’ [41]. The chasm refers to the gap between early adopters of a new technology and more pragmatic adopters; a key differentiator between these two groups is their philosophical approach to embracing novel innovations, reflected in their appetite for risk and comfort with uncertainty [9, 41].

The chasm strategy first requires identifying what is referred to as a *beachhead*: a single application area and/or industry use case where the novel innovation offers a compelling solution to problems faced in that area. Breed *et al.* [8] proposed that a compelling application for genomics is using population genomics to inform seed sourcing practices. Genomics offers critical information that is not easily attainable using other methods. For example, it can rapidly identify signals of genetic based adaptations that have the potential to increase resilience of seed stocks to future environmental change. Breed *et al.* [8] also proposed that eDNA approaches offer crucial benefits to track important ecological components and interactions during restoration (e.g., soil microbial communities; plant-pollinator communities). In addition to these application areas of genomics, beachheads can focus on a particular industry that faces a critical need to solve a problem or issue for which the novel technology is uniquely suited. For example, with respect to restoration ecology, the mining sector has increasingly regulated rehabilitation requirements that place tremendous pressure on the availability of cost-effective restoration practices [49]. The desire of scholars in our qualitative study to identify use cases where genomics offers compelling value illustrates the face validity of the ‘crossing the chasm’ model. Matching the value proposition of genomics to a critical application area allows pragmatists a compelling reason to overcome their uncertainty and move forward.

The notion of targeting a specific application area in a particular industry to gain broader market acceptance is somewhat paradoxical: why would a new technology narrow its focus rather than pursue multiple application areas and industry applications all at once? The answer lies in both the nature of word-of-mouth communications--a critical consideration for pragmatic adopters’ decision making--as well as important subtleties in how a new technology is deployed across industries and applications. Hence, a second consideration in developing a strategy for genomics to cross the chasm in restoration ecology is to explicitly consider how word-of-mouth communication might flow between areas within restoration ecology (e.g., do those who focus on seed sourcing and provenance issues interact regularly with those who focus on soil microbial communities and ecological monitoring?) as well as between different industries that face restoration pressures (e.g., do those in the mining industry share knowledge and insights with those in agriculture and/or reforestation sectors?). Another consideration is how word-of-mouth networks operate geographically, with respect to both different countries as well as types of ecosystems (rainforests vs. temperate forests vs. deserts; aquatic vs. terrestrial ecosystems).

Putting these two considerations together (selecting a beachhead and considering word-of-mouth communication) results in a strategy that, if implemented correctly, can build momentum for a new technology. A useful analogy for this strategy is to view the beachhead like a lead pin in a bowling alley: if hit properly, this lead pin will knock down adjacent pins behind it (e.g., through the word-of-mouth networks). For example, one possible “bowling alley” could start with using genomics to assess seed sourcing and transfer zones in the reclamation for degraded mining sites; then momentum could build on the one hand, to applying genomics to monitor ecological communities in those reclaimed mining sites, and on the other hand, to using genomics to inform seed sourcing and transfer zones in a related industry, such as repairing degraded agricultural ecosystems.

Finally, all new technologies require additional elements in catalysing market uptake. First, a communications strategy is required to build awareness about the value proposition that this new technology offers to the identified beachhead, as well as to educate potential adopters about how to leverage this new technology [50]. Recall that our qualitative findings highlighted the need for greater education and training with regard to what genomics is, the types of research questions it can usefully address today, and generally, building capacity, skills and knowledge. Such efforts to build awareness and capability could include workshops, education, training, and outreach, strategically delivered to a specific industry for a specific application to build momentum for communication via word-of-mouth networks to related applications and industries.

Second, funding for genomics research is equally critical. There is anecdotal evidence that some areas of restoration practice have the funding, knowledge, and motivation to use genomics, for example, to understand the impact of mining rehabilitation practices on soil microbial communities [51, 52]. Other areas of restoration suffer from insufficient funding, such as large scale restoration efforts and adequate restoration project monitoring [53], and in these cases genomics may not be easily available or a priority investment.

Third, most novel technologies require related products and services to function properly, a concept referred to as the “innovation ecosystem” [54, 55]. For example, bioinformatics and technological infrastructure are critical components for leveraging genomics. Even while attention is given to the various genomics tools and applications, equal attention must be given to properly preparing samples in the field and laboratory, and as our genomics experts noted, managing the bioinformatics challenges, and appropriately analysing and interpreting the data.

Fourth, as previously discussed, collaboration allows the benefits of bringing the power of genomics to solving restoration problems while not requiring that all members of a team be genomics experts. Rather than each person needing to bring all skills required, interdisciplinary teams are a logical solution to bring together unique and disparate skills to a project. Co-production of knowledge offers an effective approach for fostering such interdisciplinary collaboration [44].

## Conclusions

> “*This tool* [genomics] *allows us to be much more sophisticated in our understanding of ecosystem recovery following restoration*.” (Scholar 7).

Genomics offers opportunities to better understand many fundamental issues facing declining ecosystems, yet it is often a missing tool in the restoration ecologist toolbox. The goal of our study was to identify and understand the barriers slowing the uptake of genomics in restoration ecology by collecting qualitative data from semi-structured interviews and conducting an examination of research trends.

We identified varied perspectives regarding academic training and background, methodological considerations including research design and interpreting genomics data, and philosophical views regarding the benefits versus the risks of using genomics compared to traditional approaches in restoration ecology. Our interviews revealed that scholars without genomics experience feel there is perhaps a push to use the tool “*prematurely*.” Scholars with genomics experience emphatically emphasised the need to “*pump the brakes*”, to proceed cautiously in ascertaining where and how genomics can be usefully applied. The fact that categories of adopters differ with respect to perceptions and willingness to leverage novel technologies (such as genomics) is well-established in the innovation literature [9, 41]. Although our interviews offered provocative insights about genomics across various levels of experience and disciplinary expertise, our findings must be interpreted as exploratory considering our purposive sample process. For example, we did not interview junior scholars who were less experienced with genomics nor who worked only in restoration. Moreover, our interviews both preceded and prompted our subsequent examination of research trends. Hence, our scholars were not geographically representative of the recent increase in genomics research seen in Asia.

Our exploratory interviews exhibited a consensus on the need for (1) case studies to demonstrate the benefits and applications of genomics in tackling restoration problems and (2) collaboration to overcome barriers to the uptake of genomics. Evidence from our examination of research trends revealed that genomics is indeed being leveraged to address restoration issues, and in fact, its use has increased rapidly in the past few years. This increase was mostly facilitated by eDNA applications, which is by far the most widely used genomics tool, with population genomics rarely applied to restoration problems. We urge a synthesis of the 70 studies that show use of genomics to address restoration challenges.

We proposed a roadmap that explicitly considers the various aspects necessary for genomics to cross the adoption chasm. For restoration ecologists, this roadmap requires demonstrating the value of genomics in areas with a well-established ecological framework rather than applying genomics where the ecology is less well-understood; providing education, training, and outreach; ensuring funding for the research; and developing a robust set of ancillary elements (e.g., bioinformatics and computing infrastructure) to round out the necessary components of the innovation ecosystem. It would be useful to pilot the efficacy of our suggested roadmap, via studying the differential uptake in restoration across the proposed domains. With these elements in place, the likelihood of genomics to address the critical issues facing restoration will be increased.

## Acknowledgements

The authors wish to thank the interviewees for their insightful discussions that added great value to this manuscript as well as two anonymous reviewers whose comments greatly improved our manuscript. This work was supported by the Australian Research Council [grant numbers LP190100051, LP190100484, DP180100668, DP210101932].

## Declaration of interests

The authors declare no competing interests.

## Ethics

This project was done under ethics approval by University of Montana’s Ethics Review Board (approval number #144-19, for “Barriers to and Facilitators of Genomics in Ecological Restoration”) in accordance with the Code of Federal Regulations, Part 46, section 104(d).

## Supplementary information

Appendix 1. Interview guide

Appendix 2. Search strategies

Appendix 3. Inclusion/exclusion flow chart

Appendix 4. Details of the included studies

Appendix 5. Frequency of published studies from our examination of research trends by geographic region, across time

## Author Contributions

Conceptualization: JJM, PAH, MFB; Data curation: JJM, PAH, JS, MFB; Formal analysis: JJM, PAH, JS, MFB; Funding acquisition: JJM, MFB; Investigation: JJM, PAH, JS, MFB; Methodology: JJM, PAH, JS, MFB; Project administration: JJM, PAH, MFB; Resources: N/A; Software: N/A; Supervision: N/A; Validation: N/A; Visualization: PAH, JS, MFB; Writing – original draft: JJM, PAH, JS, MFB; Writing – review & editing: JJM, PAH, JS, MFB.

## Data Accessibility

The datasets supporting the examination of research trends component of this article have been uploaded as part of the Supplementary Material. The qualitative interview data are available on request.

## Notes

### Competing Interest Statement

The authors have declared no competing interest.

