## Supplementary Information for "Is the genomics ‘cart’ before the restoration ecology ‘horse’? Insights from qualitative interviews and trends from the literature" (from another version of this preprint: version 1, 2021-08-06)

[1]College of Business, Institute on Ecosystems, University of Montana, Missoula, MT, U.S.A. 59812; [2]School of Biological Sciences and Australian Research Council Training Centre for Forest Value, University of Tasmania, Hobart, Tasmania, Australia 7001; [3]School of Allied Health Science and Practice, The University of Adelaide, North Tce, Adelaide, SA, Australia 5005; [4]College of Science and Engineering, Flinders University, Bedford Park, SA, Australia 5042.

**KEYWORDS:** conservation biology, genomics, innovation, restoration ecology

**ORCIDs**: JJK 0000-0003-4938-8440, PAH 0000-0002-3502-0242, JS 0000-0002-6657-3317, MFB 0000-0001-7810-9696

**Author Contributions**: Conceptualization: JJM, PAH, MFB; Data curation: JJM, PAH, JS, MFB; Formal analysis: JJM, PAH, JS, MFB; Funding acquisition: JJM, MFB; Investigation: JJM, PAH, JS, MFB; Methodology: JJM, PAH, JS, MFB; Project administration: JJM, PAH, MFB; Resources: N/A; Software: N/A; Supervision: N/A; Validation: N/A; Visualization: PAH, JS, MFB; Writing – original draft: JJM, PAH, JS, MFB; Writing – review & editing: JJM, PAH, JS, MFB.

### Appendix 1. Interview Guide

We asked questions about the respondent's background, their understanding of genomics and how it was and/or might be used in their research field, as well as their perceptions and experience regarding barriers and facilitators.

*Background*

1. Could you describe your familiarity/experience/expertise with conservation, and restoration (e.g. how many years)?
2. How related would you consider these fields to be (e.g. subset to completely separate)?
3. How would you describe your familiarity/experience/expertise with genomics (e.g. how many years)?
   a. Could you describe what genomics is, and briefly how you learned about it?
   b. Could you provide an example use case or two of genomics in conservation and/or restoration?

*For those with genomics experience*

1. Tell us about your early experiences in bringing genomics into conservation and/or restoration.
   a. What was your motivation for adopting genomics in this way?
   b. What did you have to do to bring genomics tools/technologies into your lab?
   c. What were the early barriers? How were these barriers overcome?
   d. What were the early enablers? How did they help?
   e. What was it like (what was your experience) to secure funding? Getting published? Presenting at conferences?

*Reflective/Discipline comparisons*

1. What is your perspective about the role genomics might play in restoration ecology - the science - and/or ecosystem restoration - the practice?
2. What are your observations about the adoption rates of genomics into the two different fields? What might explain those observations?
   - History of the disciplines
   - Typical training of scientists
   - Typical tools or approaches used
   - Differences in appetite for innovation
3. Are there lessons for those in restoration ecology learnt from conservation biology, where the uptake of genomics was earlier and more rapid?

4. What advice would you give to restoration ecologists interested to increase the uptake of genomics?

### **Appendix 2. Search strategies**

#### Search strategy for *Scopus*

| Search number | Search target | Search terms |
|---|---|---|
| 1 | TITLE, ABSTRACT | Science W/3 (“conserve” OR “conservation” OR restor* OR preserv* OR “protected species” OR “protected area” OR “protected areas” OR “protected zone” OR “protected zones” OR rehabilitat* OR remediat* OR regenerat* OR rewild* OR revegetat* OR replant* OR translocat* OR reintroduc* OR re-introduc* OR re-releas* OR relocat* OR eradicate* OR biocontrol* OR bio-control* OR exclosure* OR afforestation OR reforestation) |
| 2 | AUTHOR KEYWORD | Science AND (“conserve” OR “conservation” OR restor* OR preserv* OR “protected species” OR “protected area” OR “protected areas” OR “protected zone” OR “protected zones” OR rehabilitat* OR remediat* OR regenerat* OR rewild* OR revegetat* OR replant* OR translocat* OR reintroduc* OR re-introduc* OR re-releas* OR relocat* OR eradicate* OR biocontrol* OR bio-control* OR exclosure* OR afforestation OR reforestation) |
| 3 | TITLE, ABSTRACT | biolog* W/3 (“conserve” OR “conservation” OR restor* OR preserv* OR “protected species” OR “protected area” OR “protected areas” OR “protected zone” OR “protected zones” OR rehabilitat* OR remediat* OR regenerat* OR rewild* OR revegetat* OR replant* OR translocat* OR reintroduc* OR re-introduc* OR re-releas* OR relocat* OR eradicate* OR biocontrol* OR bio-control* OR exclosure* OR afforestation OR reforestation) |
| 4 | AUTHOR KEYWORD | biolog* AND (“conserve” OR “conservation” OR restor* OR preserv* OR “protected species” OR “protected area” OR “protected areas” OR “protected zone” OR “protected zones” OR rehabilitat* OR remediat* OR regenerat* OR rewild* OR revegetat* OR replant* OR translocat* OR reintroduc* OR re-introduc* OR re-releas* OR relocat* OR eradicate* OR biocontrol* OR bio-control* OR exclosure* OR afforestation OR reforestation) |
| 5 | TITLE, ABSTRACT | ecosystem* W/3 (“conserve” OR “conservation” OR restor* OR preserv* OR “protected species” OR “protected area” OR “protected areas” OR “protected zone” OR “protected zones” OR rehabilitat* OR remediat* OR regenerat* OR rewild* OR revegetat* OR replant* OR translocat* OR reintroduc* OR re-introduc* OR re-releas* OR relocat* OR eradicate* OR biocontrol* OR bio-control* OR exclosure* OR afforestation OR reforestation) |
| 6 | AUTHOR KEYWORD | ecosystem* AND (“conserve” OR “conservation” OR restor* OR preserv* OR “protected species” OR “protected area” OR “protected areas” OR “protected zone” OR “protected zones” OR rehabilitat* OR remediat* OR regenerat* OR rewild* OR revegetat* OR replant* OR translocat* OR reintroduc* OR re-introduc* OR re-releas* OR relocat* OR eradicate* OR biocontrol* OR bio-control* OR exclosure* OR afforestation OR reforestation) |
| 7 | TITLE, ABSTRACT | ecolog* W/3 (“conserve” OR “conservation” OR restor* OR preserv* OR “protected species” OR “protected area” OR “protected areas” OR “protected zone” OR “protected zones” OR rehabilitat* OR remediat* OR regenerat* OR rewild* OR revegetat* OR replant* OR translocat* OR reintroduc* OR re-introduc* OR re-releas* OR relocat* OR eradicate* OR biocontrol* OR bio-control* OR exclosure* OR afforestation OR reforestation) |
| 8 | AUTHOR KEYWORD | ecolog* AND (“conserve” OR “conservation” OR restor* OR preserv* OR “protected species” OR “protected area” OR “protected areas” OR “protected zone” OR “protected zones” OR rehabilitat* OR remediat* OR regenerat* OR rewild* OR revegetat* OR replant* OR translocat* OR reintroduc* OR re-introduc* OR re-releas* OR relocat* OR eradicate* OR biocontrol* OR bio-control* OR exclosure* OR afforestation OR reforestation) |
| 9 | TITLE, ABSTRACT | biodivers* W/3 (“conserve” OR “conservation” OR restor* OR preserv* OR “protected species” OR “protected area” OR “protected areas” OR “protected zone” OR “protected zones” OR rehabilitat* OR remediat* OR regenerat* OR rewild* OR revegetat* OR replant* OR translocat* OR reintroduc* OR re-introduc* OR re-releas* OR relocat* OR eradicate* OR biocontrol* OR bio-control* OR exclosure* OR afforestation OR reforestation) |
| 10 | AUTHOR KEYWORD | biodivers* AND (“conserve” OR “conservation” OR restor* OR preserv* OR “protected species” OR “protected area” OR “protected areas” OR “protected zone” OR “protected zones” OR rehabilitat* OR remediat* OR regenerat* OR rewild* OR revegetat* OR replant* OR translocat* OR reintroduc* OR re-introduc* OR re-releas* OR relocat* OR eradicate* OR biocontrol* OR bio-control* OR exclosure* OR afforestation OR reforestation) |
| 11 | TITLE, ABSTRACT | habitat* W/3 (“conserve” OR “conservation” OR restor* OR preserv* OR “protected species” OR “protected area” OR “protected areas” OR “protected zone” OR “protected zones” OR rehabilitat* OR remediat* OR regenerat* OR rewild* OR revegetat* OR replant* OR translocat* OR reintroduc* OR re-introduc* OR re-releas* OR relocat* OR eradicate* OR biocontrol* OR bio-control* OR exclosure* OR afforestation OR reforestation) |

| | | |
|---|---|---|
| 12 | AUTHOR KEYWORD | habitat* AND (“conserve” OR “conservation” OR restor* OR preserv* OR “protected species” OR “protected area” OR “protected areas” OR “protected zone” OR “protected zones” OR rehabilitat* OR remediat* OR regenerat* OR rewild* OR revegetat* OR replant* OR translocat* OR reintroduc* OR re-introduc* OR re-releas* OR relocat* OR eradicate* OR biocontrol* OR bio-control* OR exclosure* OR afforestation OR reforestation) |
| 13 | | 1 OR 2 OR 3 OR 4 OR 5 OR 6 OR 7 OR 8 OR 9 OR 10 OR 11 OR 12 |
| 14 | TITLE, ABSTRACT OR AUTHOR KEYWORD | *genom* OR *exom* OR “next generation sequencing” OR “next-generation sequencing” OR “high-throughput nucleotide sequencing” OR “high throughput nucleotide sequencing” OR “massive parallel sequencing” OR meta-om* OR metaom* OR metabarcod* OR meta-barcod* OR metatranscriptom* OR meta-transcriptom* OR “high throughput sequencing” OR “high-throughput sequencing” OR “massively parallel sequencing” OR “environmental deoxyribonucleic acid” OR “eDNA” OR “environmental DNA” OR “phenome” OR “phenomes” OR “phenomics” OR “phenomic” |
| 15 | | 13 AND 14 |

### Search strategy for *Web of Science Core Collection*

| Search number | Search target | Search terms |
|---|---|---|
| 1 | TITLE, ABSTRACT | Science NEAR/3 (“conserve” OR “conservation” OR restor* OR preserv* OR “protected species” OR “protected area” OR “protected areas” OR “protected zone” OR “protected zones” OR rehabilitat* OR remediat* OR regenerat* OR rewild* OR revegetat* OR replant* OR translocat* OR reintroduc* OR re-introduc* OR re-releas* OR relocat* OR eradicate* OR biocontrol* OR bio-control* OR exclosure* OR afforestation OR reforestation) |
| 2 | AUTHOR KEYWORD | Science AND (“conserve” OR “conservation” OR restor* OR preserv* OR “protected species” OR “protected area” OR “protected areas” OR “protected zone” OR “protected zones” OR rehabilitat* OR remediat* OR regenerat* OR rewild* OR revegetat* OR replant* OR translocat* OR reintroduc* OR re-introduc* OR re-releas* OR relocat* OR eradicate* OR biocontrol* OR bio-control* OR exclosure* OR afforestation OR reforestation) |
| 3 | TITLE, ABSTRACT | biolog* NEAR/3 (“conserve” OR “conservation” OR restor* OR preserv* OR “protected species” OR “protected area” OR “protected areas” OR “protected zone” OR “protected zones” OR rehabilitat* OR remediat* OR regenerat* OR rewild* OR revegetat* OR replant* OR translocat* OR reintroduc* OR re-introduc* OR re-releas* OR relocat* OR eradicate* OR biocontrol* OR bio-control* OR exclosure* OR afforestation OR reforestation) |
| 4 | AUTHOR KEYWORD | biolog* AND (“conserve” OR “conservation” OR restor* OR preserv* OR “protected species” OR “protected area” OR “protected areas” OR “protected zone” OR “protected zones” OR rehabilitat* OR remediat* OR regenerat* OR rewild* OR revegetat* OR replant* OR translocat* OR reintroduc* OR re-introduc* OR re-releas* OR relocat* OR eradicate* OR biocontrol* OR bio-control* OR exclosure* OR afforestation OR reforestation) |
| 5 | TITLE, ABSTRACT | ecosystem* NEAR/3 (“conserve” OR “conservation” OR restor* OR preserv* OR “protected species” OR “protected area” OR “protected areas” OR “protected zone” OR “protected zones” OR rehabilitat* OR remediat* OR regenerat* OR rewild* OR revegetat* OR replant* OR translocat* OR reintroduc* OR re-introduc* OR re-releas* OR relocat* OR eradicate* OR biocontrol* OR bio-control* OR exclosure* OR afforestation OR reforestation) |
| 6 | AUTHOR KEYWORD | ecosystem* AND (“conserve” OR “conservation” OR restor* OR preserv* OR “protected species” OR “protected area” OR “protected areas” OR “protected zone” OR “protected zones” OR rehabilitat* OR remediat* OR regenerat* OR rewild* OR revegetat* OR replant* OR translocat* OR reintroduc* OR re-introduc* OR re-releas* OR relocat* OR eradicate* OR biocontrol* OR bio-control* OR exclosure* OR afforestation OR reforestation) |
| 7 | TITLE, ABSTRACT | ecolog* NEAR/3 (“conserve” OR “conservation” OR restor* OR preserv* OR “protected species” OR “protected area” OR “protected areas” OR “protected zone” OR “protected zones” OR rehabilitat* OR remediat* OR regenerat* OR rewild* OR revegetat* OR replant* OR translocat* OR reintroduc* OR re-introduc* OR re-releas* OR relocat* OR eradicate* OR biocontrol* OR bio-control* OR exclosure* OR afforestation OR reforestation) |
| 8 | AUTHOR KEYWORD | ecolog* AND (“conserve” OR “conservation” OR restor* OR preserv* OR “protected species” OR “protected area” OR “protected areas” OR “protected zone” OR “protected zones” OR rehabilitat* OR remediat* OR regenerat* OR rewild* OR revegetat* OR replant* OR translocat* OR reintroduc* OR re-introduc* OR re-releas* OR relocat* OR eradicate* OR biocontrol* OR bio-control* OR exclosure* OR afforestation OR reforestation) |

| | | |
|---|---|---|
| 9 | TITLE, ABSTRACT | biodivers* NEAR/3 (“conserve” OR “conservation” OR restor* OR preserv* OR “protected species” OR “protected area” OR “protected areas” OR “protected zone” OR “protected zones” OR rehabilitat* OR remediat* OR regenerat* OR rewild* OR revegetat* OR replant* OR translocat* OR reintroduc* OR re-introduc* OR re-releas* OR relocat* OR eradicate* OR biocontrol* OR bio-control* OR exclosure* OR afforestation OR reforestation) |
| 10 | AUTHOR KEYWORD | biodivers* AND (“conserve” OR “conservation” OR restor* OR preserv* OR “protected species” OR “protected area” OR “protected areas” OR “protected zone” OR “protected zones” OR rehabilitat* OR remediat* OR regenerat* OR rewild* OR revegetat* OR replant* OR translocat* OR reintroduc* OR re-introduc* OR re-releas* OR relocat* OR eradicate* OR biocontrol* OR bio-control* OR exclosure* OR afforestation OR reforestation) |
| 11 | TITLE, ABSTRACT | habitat* NEAR/3 (“conserve” OR “conservation” OR restor* OR preserv* OR “protected species” OR “protected area” OR “protected areas” OR “protected zone” OR “protected zones” OR rehabilitat* OR remediat* OR regenerat* OR rewild* OR revegetat* OR replant* OR translocat* OR reintroduc* OR re-introduc* OR re-releas* OR relocat* OR eradicate* OR biocontrol* OR bio-control* OR exclosure* OR afforestation OR reforestation) |
| 12 | AUTHOR KEYWORD | habitat* AND (“conserve” OR “conservation” OR restor* OR preserv* OR “protected species” OR “protected area” OR “protected areas” OR “protected zone” OR “protected zones” OR rehabilitat* OR remediat* OR regenerat* OR rewild* OR revegetat* OR replant* OR translocat* OR reintroduc* OR re-introduc* OR re-releas* OR relocat* OR eradicate* OR biocontrol* OR bio-control* OR exclosure* OR afforestation OR reforestation) |
| 13 | | 1 OR 2 OR 3 OR 4 OR 5 OR 6 OR 7 OR 8 OR 9 OR 10 OR 11 OR 12 |
| 14 | TITLE | *genom* OR *exom* OR “next generation sequencing” OR “next-generation sequencing” OR “high-throughput nucleotide sequencing” OR “high throughput nucleotide sequencing” OR “massive parallel sequencing” OR meta-om* OR metaom* OR metabarcod* OR meta-barcod* OR metatranscriptom* OR meta-transcriptom* OR “high throughput sequencing” OR “high-throughput sequencing” OR “massively parallel sequencing” OR “environmental deoxyribonucleic acid” OR “eDNA” OR “environmental DNA” OR “phenome” OR “phenomes” OR “phenomics” OR “phenomic” |
| 15 | ABSTRACT, AUTHOR KEYWORD | meta-genom* OR metagenom* OR genom* OR whole-genom* OR whole-exom* OR exom* OR “next generation sequencing” OR “next-generation sequencing” OR “high-throughput nucleotide sequencing” OR “high throughput nucleotide sequencing” OR “massive parallel sequencing” OR meta-om* OR metaom* OR metabarcod* OR meta-barcod* OR metatranscriptom* OR meta-transcriptom* OR “high throughput sequencing” OR “high-throughput sequencing” OR “massively parallel sequencing” OR-“environmental deoxyribonucleic acid” OR “eDNA” OR “environmental DNA” OR “phenome” OR “phenomes” OR “phenomics” OR “phenomic” |
| 16 | | 14 OR 15 |
| 17 | | 13 AND 16 |

**Appendix 3. Inclusion/exclusion flow chart**

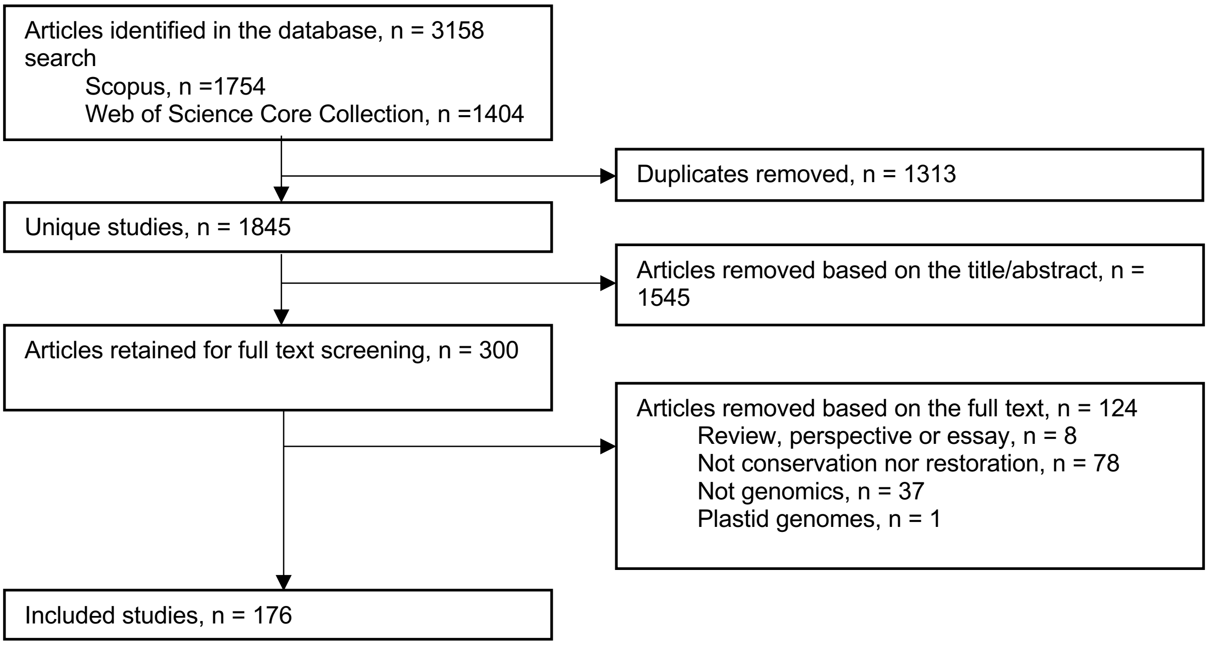

### Appendix 4. Details of the included studies

| Reference | Country of data collection | Country of author affiliation(s) | Year(s) of data collection | Restoration or conservation (author) | Restoration or conservation (reviewer) | Type of genomics |
|---|---|---|---|---|---|---|
| Abrams, J.F., Hörig, L.A., Brozovic, R., Axtner, J., Crampton-Platt, A., Mohamed, A., Wong, S.T., Sollmann, R., Yu, D.W. & Wilting, A. 2019 Shifting up a gear with iDNA: from mammal detection events to standardised surveys. Journal of Applied Ecology 56, 1637-1648. (doi:10.1111/1365-2664.13411). | Malaysia | Germany, USA, UK, China | 2015 | Conservation | Conservation | Environmental DNA |
| Ahrendsen, D.L., Aust, S.K. & Kellar, P.R. 2016 Biodiversity assessment using next-generation sequencing: comparison of phylogenetic and functional diversity between Nebraska grasslands. Plant Systematics and Evolution 302, 89-108. (doi:10.1007/s00606-015-1246-6). | USA | USA | NR | Restoration | Restoration | Environmental DNA |
| Allen, J.L., McKenzie, S.K., Sleith, R.S. & Alter, S.E. 2018 First genome-wide analysis of the endangered, endemic lichen Cetradonia linearis reveals isolation by distance and strong population structure. American Journal of Botany 105, 1556-1567. (doi:10.1002/ajb2.1150). | USA | USA | NR | Conservation | Conservation | Population genomics |
| Andersen, L.H., Sunde, P., Pellegrino, I., Loeschcke, V. & Pertoldi, C. 2017 Using population viability analysis, genomics, and habitat suitability to forecast future population patterns of Little Owl Athene noctua across Europe. Ecology and Evolution 7, 10987-11001. (doi:10.1002/ece3.3629). | “Europe” | Denmark, Italy | NR | Conservation | Conservation | Population genomics |
| Ando, H., Setsuko, S., Horikoshi, K., Suzuki, H., Umehara, S., Inoue-Murayama, M. & Isagi, Y. 2013 Diet analysis by next-generation sequencing indicates the frequent consumption of introduced plants by the critically endangered red-headed wood pigeon (Columba janthina nitens) in oceanic island habitats. Ecology and Evolution 3, 4057-4069. (doi:10.1002/ece3.773). | Japan | Japan | 2009-2011 | Conservation | Conservation | Environmental DNA |
| Angeloni, F., Wagemaker, C.A.M., Jetten, M.S.M., Op den Camp, H.J.M., Janssen-Megens, E.M., Francoijs, K.-J., Stunnenberg, H.G. & Ouborg, N.J. 2011 De novo transcriptome characterization and development of genomic tools for Scabiosa columbaria L. using next-generation sequencing techniques. Molecular Ecology Resources 11, 662-674. (doi:10.1111/j.1755-0998.2011.02990.x). | UK | Netherlands | NR | Conservation | Conservation | Population genomics |
| Anthony, M.A., Stinson, K.A., Trautwig, A.N., Coates-Connor, E. & Frey, S.D. 2019 Fungal communities do not recover after removing invasive Alliaria petiolata (garlic mustard). Biological Invasions 21, 2085-2099. (doi:10.1007/s10530-019-02031-8). | USA | USA | 2013-2017 | Restoration | Restoration | Environmental DNA |

| | | | | | | |
|---|---|---|---|---|---|---|
| Arrizabalaga-Escudero, A., Garin, I., García-Mudarra, J.L., Alberdi, A., Aihartza, J. & Goiti, U. 2015 Trophic requirements beyond foraging habitats: the importance of prey source habitats in bat conservation. Biological Conservation 191, 512-519. (doi:10.1016/j.biocon.2015.07.043). | Spain | Spain, The Basque Country | 2012 | Conservation | Conservation | Environmental DNA |
| Axtner, J., Crampton-Platt, A., Hörig, L.A., Mohamed, A., Xu, C.C.Y., Yu, D.W. & Wilting, A. 2019 An efficient and robust laboratory workflow and tetrapod database for larger scale environmental DNA studies. GigaScience 8, 1-17. (doi:10.1093/gigascience/giz029). | Malaysia | Germany, China, Netherlands, Canada, UK | Database up until 2015 | Conservation | Conservation | Environmental DNA |
| Bálint, M., Nowak, C., Márton, O., Pauls, S.U., Wittwer, C., Aramayo, J.L., Schulze, A., Chambert, T., Cocchiararo, B. & Jansen, M. 2018 Accuracy, limitations and cost efficiency of eDNA-based community survey in tropical frogs. Molecular Ecology Resources 18, 1415-1426. (doi:10.1111/1755-0998.12934). | Bolivia | Germany, Hungary, Bolivia, USA | 2014 | Conservation | Conservation | Environmental DNA |
| Baloğlu, B., Clews, E. & Meier, R. 2018 NGS barcoding reveals high resistance of a hyperdiverse chironomid (Diptera) swamp fauna against invasion from adjacent freshwater reservoirs. Frontiers in Zoology 15, 31. (doi:10.1186/s12983-018-0276-7). | Singapore | Singapore | 2013-2014 | Conservation | Conservation | Environmental DNA |
| Barboza, A.D.M., Pylro, V.S., Jacques, R.J.S., Gubiani, P.I., de Quadros, F.L.F., da Trindade, J.K., Triplett, E.W. & Foesch, L. 2018 Seasonal dynamics alter taxonomical and functional microbial profiles in Pampa biome soils under natural grasslands. PeerJ 6, e4991. (doi:10.7717/peerj.4991). | Brazil | Brazil, USA | 2014-2015 | Conservation | Conservation | Environmental DNA |
| Bellemain, E., Patricio, H., Gray, T., Guegan, F., Valentini, A., Miaud, C. & Dejean, T. 2016 Trails of river monsters: Detecting critically endangered Mekong giant catfish Pangasianodon gigas using environmental DNA. Global Ecology and Conservation 7, 148-156. (doi:10.1016/j.gecco.2016.06.007). | Thailand | France, Australia, Cambodia, Lao Democratic People's Republic | 2015 | Conservation | Conservation | Environmental DNA |
| Bessey, C., Jarman, S.N., Stat, M., Rohner, C.A., Bunce, M., Koziol, A., Power, M., Rambahiniarison, J.M., Richardson, A.J. & Berry, O. 2019 DNA metabarcoding assays reveal a diverse prey assemblage for Mobula rays in the Bohol Sea, Philippines. Ecology and Evolution 9, 2459-2474. (doi:10.1002/ece3.4858). | Philippines | Australia, USA, Philippines | 2013-2015 | Conservation | Conservation | Environmental DNA |
| Bi, K., Linderoth, T., Vanderpool, D., Good, J.M., Nielsen, R. & Moritz, C. 2013 Unlocking the vault: next-generation museum population genomics. Molecular Ecology 22, 6018-6032. (doi:10.1111/mec.12516). | USA | USA, Australia | 1915-2012 | Conservation | Conservation | Population genomics |

| | | | | | | |
|---|---|---|---|---|---|---|
| Biggs, J., Ewald, N., Valentini, A., Gaboriaud, C., Dejean, T., Griffiths, R.A., Foster, J., Wilkinson, J.W., Arnell, A., Brotherton, P., et al. 2015 Using eDNA to develop a national citizen science-based monitoring programme for the great crested newt (Triturus cristatus). Biological Conservation 183, 19-28. (doi:10.1016/j.biocon.2014.11.029). | UK | UK, France | 2013 | Conservation | Conservation | Environmental DNA |
| Binks, R.M., Byrne, M., McMahon, K., Pitt, G., Murray, K. & Evans, R.D. 2019 Habitat discontinuities form strong barriers to gene flow among mangrove populations, despite the capacity for long-distance dispersal. Diversity and Distributions 25, 298-309. (doi:10.1111/ddi.12851). | Australia | Australia | NR | Conservation | Conservation | Population genomics |
| Boeraeve, M., Honnay, O., Mullens, N., Vandekerkhove, K., De Keersmaeker, L., Thomaes, A. & Jacquemyn, H. 2018 The impact of spatial isolation and local habitat conditions on colonization of recent forest stands by ectomycorrhizal fungi. Forest Ecology and Management 429, 84-92. (doi:10.1016/j.foreco.2018.06.043). | Belgium | Belgium | NR | Restoration | Restoration | Environmental DNA |
| Boessenkool, S., McGlynn, G., Epp, L.S., Taylor, D., Pimentel, M., Gizaw, A., Nemomissa, S., Brochmann, C. & Popp, M. 2014 Use of ancient sedimentary DNA as a novel conservation tool for high-altitude tropical biodiversity. Conservation Biology 28, 446-455. (doi:10.1111/cobi.12195). | Uganda, Rwanda | Norway, Ireland, Singapore, Ethiopia | 2018 | Both | Both | Environmental DNA |
| Boscari, E., Abbiati, M., Badalamenti, F., Bavestrello, G., Benedetti-Cecchi, L., Cannas, R., Cau, A., Cerrano, C., Chimienti, G., Costantini, F., et al. 2019 A population genomics insight by 2b-RAD reveals populations' uniqueness along the Italian coastline in Leptopsammia pruvoti (Scleractinia, Dendrophylliidae). Diversity and Distributions 25, 1101-1117. (doi:10.1111/ddi.12918). | Italy | Italy | 2014-2015 | Conservation | Conservation | Population genomics |
| Brauer, C.J., Hammer, M.P. & Beheregaray, L.B. 2016 Riverscape genomics of a threatened fish across a hydroclimatically heterogeneous river basin. Molecular Ecology 25, 5093-5113. (doi:10.1111/mec.13830). ] | Australia | Australia | 2000-2013 | Conservation | Conservation | Population genomics |
| Brozio, S., Manson, C., Gourevitch, E., Burns, T.J., Greener, M.S., Downie, J.D. & Hoskinsson, P.A. 2017 Development and application of an eDNA method to detect the critically endangered Trinidad golden tree frog (Phytotriades auratus) in Bromeliad Phytotelmata. PLoS One 12, e0170619. (doi:10.1371/journal.pone.0170619). | Trinidad | UK | 2015 | Conservation | Conservation | Environmental DNA |
| Brunbjerg, A.K., Bruun, H.H., Dalby, L., Fløjgaard, C., Frøslev, T.G., Høye, T.T., Goldberg, I., Læssøe, T., Hansen, M.D.D., Brøndum, L., et al. 2018 Vascular plant species richness and bioindication predict multi-taxon species richness. Methods in Ecology and Evolution 9, 2372-2382. (doi:10.1111/2041-210X.13087). | Denmark | Denmark | NR | Conservation | Both | Environmental DNA |

| | | | | | | |
|---|---|---|---|---|---|---|
| Brüniche-Olsen, A., Austin, J.J., Jones, M.E., Holland, B.R. & Burridge, C.P. 2016 Detecting selection on temporal and spatial scales: a genomic time-series assessment of selective responses to devil facial tumor disease PLoS One 11, e0147875. (doi:10.1371/journal.pone.0147875). | Australia | Australia | 1999-2013 | Conservation | Conservation | Population genomics |
| Brüniche-Olsen, A., Kellner, K.F. & DeWoody, J.A. 2019 Island area, body size and demographic history shape genomic diversity in Darwin's finches and related tanagers. Molecular Ecology 28, 4914-4925. (doi:10.1111/mec.15266). | Ecuador | USA | 1988-2001 | Conservation | Both | Population genomics |
| Burgar, J.M., Murray, D.C., Craig, M.D., Haile, J., Houston, J., Stokes, V. & Bunce, M. 2014 Who's for dinner? High-throughput sequencing reveals bat dietary differentiation in a biodiversity hotspot where prey taxonomy in largely undescribed. Molecular Ecology 23, 3605-3617. (doi:10.1111/mec.12531). | Australia | Australia | 2010-2011 | Conservation | Conservation | Environmental DNA |
| Burgess, T.I., White, D., McDougall, K.M., Garnas, J., Dunstan, W.A., Català, S., Carnegie, A.J., Worboys, S., Cahill, D., Vettraino, A.-M., et al. 2017 Distribution and diversity of Phytophthora across Australia. Pacific Conservation Biology 23, 150-162. (doi:10.1071/PC16032). | Australia | Australia, Italy, Spain, South Africa | 2013-2015 | Conservation | Conservation | Environmental DNA |
| Callicrate, T., Dikow, R., Thomas, J.W., Mullikin, J.C., Jarvis, E.D., Fleischer, R.C. & NISC Comparative Sequencing Program. 2014 Genomic resources for the endangered Hawaiian honeycreepers. BMC Genomics 15, 1098. (doi:10.1186/1471-2164-15-1098). | USA | USA | 2002 | Conservation | Conservation | Comparative genomics |
| Castañeda, L.E. & Barbosa, O. 2017 Metagenomic analysis exploring taxonomic and functional diversity of soil microbial communities in Chilean vineyards and surrounding native forests. PeerJ 5, e3098. (doi:10.7717/peerj.3098). | Chile | Chile | 2012 | Conservation | Both | Environmental DNA |
| Chen, J., Mo, L., Zhang, Z., Nan, J., Xu, D., Chao, L., Zhang, X. & Bao, Y. 2020 Evaluation of the ecological restoration of a coal mine dump by exploring the characteristics of microbial communities. Applied Soil Ecology 147, 103430. (doi:10.1016/j.apsoil.2019.103430). | China | China | 2018 | Restoration | Restoration | Environmental DNA |
| Chen, N., Cosgrove, E.J., Bowman, R., Fitzpatrick, J.W. & Clark, A.G. 2016 Genomic consequences of population decline in the endangered Florida scrub-jay. Current Biology 26, 2974-2979. (doi:10.1016/j.cub.2016.08.062). | USA | USA | 1995-2013 | Conservation | Conservation | Population genomics |

| | | | | | | |
|---|---|---|---|---|---|---|
| Çilingir, F.G., Rheindt, F.E., Garg, K.M., Platt, K., Platt, S.G. & Bickford, D.P. 2017 Conservation genomics of the endangered Burmese roofed turtle. Conservation Biology 31, 1469-1476. (doi:10.1111/cobi.12921). | Myanmar | Singapore, Myanmar, USA | 2006-2011 | Conservation | Conservation | Population genomics |
| Cilleros, K., Valentini, A., Allard, L., Dejean, T., Etienne, R., Grenouillet, G., Iribar, A., Taberlet, P., Vigouroux, R. & Brosse, S. 2019 Unlocking biodiversity and conservation studies in high-diversity environments using environmental DNA (eDNA): a test with Guianese freshwater fishes. Molecular Ecology Resources 19, 27-46. (doi:10.1111/1755-0998.12900). | French Guiana | France, French Guiana | 2010-2015 | Conservation | Conservation | Environmental DNA |
| Cordeiro, E.M.G., Macrini, C.M., Sujii, P.S., Schwarcz, K.D., Pinheiro, J.B., Rodrigues, R.R., Brancalion, P.H.S. & Zucchi, M.I. 2019 Diversity, genetic structure, and population genomics of the tropical tree Centrolobium tomentosum in remnant and restored Atlantic forests. Conservation Genetics 20, 1073-1085. (doi:10.1007/s10592-019-01195-z). | Brazil | Brazil | 2013-2014 | Both | Both | Population genomics |
| Cristofari, R., Plaza, P., Fernández, C.E., Trucchi, E., Gouin, N., Le Bohec, C., Zavalaga, C., Alfaro-Shigueto, J. & Luna-Jorquera, G. 2019 Unexpected population fragmentation in an endangered seabird: the case of the Peruvian diving-petrel. Scientific Reports 9, 2021. (doi:10.1038/s41598-019-38682-9). | Chile | Finland, France, Monaco, Norway, Chile, Austria, Italy, Peru | 2012 | Conservation | Conservation | Population genomics |
| Cruz, M.V., Mori, G.M., Signori-Müller, C., da Silva, C.C., Oh, D.-H., Dassanayake, M., Zucchi, M.I., Oliveira, R.S. & de Souza, A.P. 2019 Local adaptation of a dominant coastal tree to freshwater availability and solar radiation suggested by genomic and ecophysiological approaches. Scientific Reports 9, 19936. (doi:10.1038/s41598-019-56469-w). | "South America" | Brazil, USA | 2014 | Conservation | Conservation | Population genomics |
| De Barba, M., Miquel, C., Boyer, F., Mercier, C., Rioux, D., Coissac, E. & Taberlet, P. 2014 DNA metabarcoding multiplexing and validation of data accuracy for diet assessment: application to omnivorous diet. Molecular Ecology Resources 14, 306-323. (doi:10.1111/1755-0998.12188). | Italy | Moscow | 2002-2008 | Conservation | Conservation | Environmental DNA |
| de Villemereuil, P., Rutschmann, A., Lee, K.D., Ewen, J.G., Brekke, P. & Santure, A.W. 2019 Little adaptive potential in a threatened passerine bird. Current Biology 29, 889-894. (doi:10.1016/j.cub.2019.01.072). | New Zealand | France, UK | 2004-2015 | Conservation | Conservation | Population genomics |
| Devitt, T.J., Wright, A.M., Cannatella, D.C. & Hillis, D.M. 2019 Species delimitation in endangered groundwater salamanders: implications for aquifer management and biodiversity conservation. Proceedings of the National Academy of Science 116, 2624-2633. (doi:10.1073/pnas.1815014116). | USA | USA | NR | Conservation | Conservation | Population genomics |

| | | | | | | |
|---|---|---|---|---|---|---|
| DeWoody, J.A., Fernandez, N.B., Brüniche-Olsen, A., Antonides, J.D., Doyle, J.M., San Miguel, P., Westerman, R., Vertyankin, V.V., Godard-Codding, C.A.J. & Bickham, J.W. 2017 Characterization of the grey whale Eschrichtius robustus genome and a genotyping array based on single-nucleotide polymorphisms in candidate genes. The Biological Bulletin 232, 186-197. (doi:10.1086/693483). | "Pacific" | USA, Russia | NR | Conservation | Conservation | Population genomics |
| Dittberner, H., Becker, C., Jiao, W.-B., Schneeberger, K., Hölzel, N., Tellier, A. & de Meaux, J. 2019 Strengths and potential pitfalls of hay transfer for ecological restoration revealed by RAD-seq analysis in floodplain Arabis species. Molecular Ecology 28, 3887-3901. (doi:10.1111/mec.15194). | Germany | Germany | 1997-2016 | Restoration | Restoration | Population genomics & environmental DNA |
| Dobrynin, P., Liu, S., Tamazian, G., Xiong, Z., Yurchenko, A.A., Krasheninnikova, K., Kliver, S., Schmidt-Küntzel, A., Koepfli, K.-P., Johnson, W., et al. 2015 Genomic legacy of the African cheetah, Acinonyx jubatus. Genome Biology 16, 277. (doi:10.1186/s13059-015-0837-4). | Namibia, Tanzania | Russia, China, Namibia, USA, Spain, Portugal, Puerto Rico, Brazil, Korea, Denmark, Saudi Arabia | NR | Conservation | Conservation | Population genomics |
| Docherty, K.M. & Gutknecht, J.L.M. 2019 Soil microbial restoration strategies for promoting climate-ready prairie ecosystems. Ecological Applications 29, e01858. (doi:10.1002/eap.1858). | USA | USA | 2015 | Restoration | Restoration | Environmental DNA |
| Drinkwater, R., Bærholm Schnell, I., Bohmann, K., Bernard, H., Veron, G., Clare, E., Gilbert, M.T.P. & Rossiter, S.J. 2019 Using metabarcoding to compare the suitability of two blood-feeding leech species for sampling mammalian diversity in North Borneo. Molecular Ecology Resources 19, 105-117. (doi:10.1111/1755-0998.12943). | Malaysia | UK, Denmark, Malaysia, France, Norway | 2015 | Conservation | Conservation | Environmental DNA |
| Drury, C., Schopmeyer, S., Goergen, E., Bartels, E., Nedimyer, K., Johnson, M., Maxwell, K., Galvan, V., Manfrino, C. & Lirman, D. 2017 Genomic patterns in Acropora cervicornis show extensive population structure and variable genetic diversity Ecology and Evolution 7, 6188-6200. (doi:10.1002/ece3.3184). | USA | Dominican Republic, USA, Cayman Islands | 2012-2015 | Conservation | Conservation | Population genomics |
| Dupuis, J.R., Oliver, J.C., Brunet, B.M.T., Longcore, T., Johnson, J.J. & Sperling, F.A.H. 2018 Genomic data indicate ubiquitous evolutionary distinctiveness among populations of California metalmark butterflies. Conservation Genetics 19, 1097-1108. (doi:10.1007/s10592-018-1081-8). | USA | USA, Canada | 1995-2014 | Conservation | Conservation | Population genomics |

| | | | | | | |
|---|---|---|---|---|---|---|
| Dupuis, J.R., Peigler, R.S., Geib, S.M. & Rubinoff, D. 2018 Phylogenomics supports incongruence between ecological specialization and taxonomy in a charismatic clade of buck moths. Molecular Ecology 27, 4417-4429. (doi:10.1111/mec.14883). | USA | USA | 1997-2015 | Conservation | Conservation | Population genomics |
| Eaton, W.D., Shokralla, S., McGee, K.M. & Hajibabaei, M. 2017 Using metagenomics to show the efficacy of forest restoration in the New Jersey Pine Barrens. Genome 60, 825-836. (doi:10.1139/gen-2015-0199). | USA | USA, Canada, Egypt | 2011 | Restoration | Restoration | Environmental DNA |
| Ejrnæs, R., Frøslev, T.G., Høye, T.T., Kjøller, R., Oddershede, A., Brunbjerg, A.K., Hansen, A.J. & Bruun, H.H. 2018 Uniquity: A general metric for biotic uniqueness of sites. Biological Conservation 225, 98-105. (doi:10.1016/j.biocon.2018.06.034). | Denmark | Denmark | 2014-2016 | Conservation | Both | Environmental DNA |
| Ekblom, R., Brechlin, B., Persson, J., Smeds, L., Johansson, M., Magnusson, J., Flagstad, Ø. & Ellegren, H. 2018 Genome sequencing and conservation genomics in the Scandinavian wolverine population Conservation Biology 32, 1301-1312. (doi:10.1111/cobi.13157). | Sweden | Sweden, Norway | 2005-2010 | Conservation | Conservation | Population genomics |
| Esnaola, A., Arrizabalaga-Escudero, A., González-Esteban, J., Elosegi, A. & Aihartza, J. 2018 Determining diet from faeces: Selection of metabarcoding primers for the insectivore Pyrenean desman (Galemys pyrenaicus). PLoS One 13, e0208986. (doi:10.1371/journal.pone.0208986). | Spain | Spain | 2015-2016 | Conservation | Conservation | Environmental DNA |
| Ezeokoli, O.T., Bezuidenhout, C.C., Maboeta, M.S., Khasa, D.P. & Adeleke, R.A. 2020 Structural and functional differentiation of bacterial communities in post-coal mining reclamation soils of South Africa: bioindicators of soil ecosystem restoration. Scientific Reports 10, 1759. (doi:10.1038/s41598-020-58576-5). | South Africa | South Africa, Canada | 2016 | Restoration | Restoration | Environmental DNA |
| Feng, S., Fang, Z., Barnett, R., Li, C., Han, S., Kuhlwilm, M., Zhou, L., Pan, H., Deng, Y., Chen, G., et al. 2019 The genomic footprints of the fall and recovery of the crested ibis. Current Biology 29, 340-349. (doi:10.1016/j.cub.2018.12.008). | China, Korea, Japan, Russia | China, Denmark, UK, Spain, Austria, Germany, Norway | 1841-1922 | Conservation | Conservation | Population genomics |
| Fernandes, K., van der Heyde, M., Coghlan, M., Wardell-Johnson, G., Bunce, M., Harris, R. & Nevill. 2019 Invertebrate DNA metabarcoding reveals changes in communities across mine site restoration chronosequences. Restoration Ecology 27, 1177-1186. (doi:10.1111/rec.12976). | Australia | Australia | 2017 | Restoration | Restoration | Environmental DNA |
| Feutry, P., Berry, O., Kyne, P.M., Pillans, R.D., Hillary, R.M., Grewe, P.M., Marthick, J.R., Johnson, G., Gunasekera, R.M., Bax, N.J., et al. 2017 Inferring contemporary and historical | Australia | Australia | 2012-2014 | Conservation | Conservation | Population genomics |

| | | | | | | |
|---|---|---|---|---|---|---|
| genetic connectivity from juveniles. Molecular Ecology 26, 444-456. (doi:10.1111/mec.13929). | | | | | | |
| Ficetola, G.F., Miaud, C., Pompanon, F. & Taberlet, P. 2008 Species detection using environmental DNA from water samples. Biology Letters 4, 423-425. (doi:10.1098/rsbl.2008.0118). | France | France | NR | Conservation | Both | Environmental DNA |
| Fløjgaard, C., De Barba, M., Taberlet, P. & Ejrnæs, R. 2017 Body condition, diet and ecosystem function of red deer (Cervus elaphus) in a fenced nature reserve. Global Ecology and Conservation 11, 312-323. (doi:10.1016/j.gecco.2017.07.003). | Denmark | Denmark, France | 2013-2014 | Conservation | Conservation | Environmental DNA |
| Garbe, J.R., Prakapenka, D., Tan, C. & Da, Y. 2016 Genomic inbreeding and relatedness in wild panda populations. PLoS One 11, e0160496. (doi:10.1371/journal.pone.0160496). | China | USA, China | NR | Conservation | Conservation | Population genomics |
| Gellie, N.J.C., Mills, J.G., Breed, M.F. & Lowe, A.J. 2017 Revegetation rewilds the soil bacterial microbiome of an old field. Molecular Ecology 26, 2895-2904. (doi:10.1111/mec.14081). | Australia | Australia | 2015 | Restoration | Restoration | Environmental DNA |
| Geml, J., Gravendeel, B., van der Gaag, K., Neilen, M., Lammers, Y., Raes, N., Semenova, T.A., de Knijff, P. & Noordeloos, M.E. 2014 The contribution of DNA metabarcoding to fungal conservation: diversity assessment, habitat partitioning and mapping red-listed fungi in protected coastal Salix repens communities in the Netherlands. PLoS One 9, e99852. (doi:10.1371/journal.pone.0099852). | Netherlands | Netherlands | 2010 | Conservation | Conservation | Environmental DNA |
| Grange, Z.L., Biggs, P.J., Rose, S.P., Gartrell, B.D., Nelson, N.J. & French, N.P. 2017 Genomic epidemiology and management of Salmonella in island ecosystems used for takahe conservation. Microbial Ecology 74, 735-744. (doi:10.1007/s00248-017-0959-1). | New Zealand | New Zealand, USA | 2011-2013 | Conservation | Conservation | Comparative genomics |
| Grange, Z.L., Gartrell, B.D., Biggs, P.J., Nelson, N.J., Anderson, M. & French, N.P. 2016 Microbial genomics of a host-assocated commensal bacterium in fragmented populations of endangered takahe. Microbial Ecology 71, 1020-1029. (doi:10.1007/s00248-015-0721-5). | New Zealand | New Zealand | 2011-2013 | Conservation | Conservation | Comparative genomics |
| Gueuning, M., Ganser, D., Blaser, S., Albrecht, M., Knop, E., Praz, C. & Frey, J.E. 2019 Evaluating next-generation sequencing (NGS) methods for routine monitoring of wild bees: Metabarcoding, mitogenomics or NGS barcoding. Molecular Ecology Resources 19, 847-862. (doi:10.1111/1755-0998.13013). | Switzerland | Switzerland | 2017 | Conservation | Both | Environmental DNA |

| | | | | | | |
|---|---|---|---|---|---|---|
| Guo, W., Mishra, S., Wang, C., Zhang, H., Ning, R., Kong, F., Zeng, B., Zhao, J. & Li, Y. 2019 Comparative Study of Gut Microbiota in Wild and Captive Giant Pandas (Ailuropoda melanoleuca). Genes (Basel) 10, 827. (doi:10.3390/genes10100827). | China | China, USA | 2013-2015 | Conservation | Conservation | Environmental DNA |
| Guo, Y., Chen, X., Wu, Y., Zhang, L., Cheng, J., Wei, G. & Lin, Y. 2018 Natural revegetation of a semiarid habitat alters taxonomic and functional diversity of soil microbial communities. Science of the Total Environment 635, 598-606. (doi:10.1016/j.scitotenv.2018.04.171). | China | China | 2014 | Restoration | Restoration | Environmental DNA |
| Guyton, J.A., Pansu, J., Hutchinson, M.C., Kartzinel, T.R., Potter, A.B., Coverdale, T.C., Daskin, J.H., da Conceição, A.G., Peel, M.J.S., Stalmans, M.E., et al. 2020 Trophic rewilding revives biotic resistance to shrub invasion. Nature Ecology & Evolution 4, 712-724. (doi:10.1038/s41559-019-1068-y). | Mozambique | USA, France, Australia, Mozambique, South Africa | 2013-2018 | Conservation | Conservation | Environmental DNA |
| Halifu, S., Deng, X., Song, X., An, Y. & Song, R. 2019 Effects of sphaeropsis blight on rhizosphere soil bacterial community structure and soil physicochemical properties of Pinus sylvestris var. mongolica in Zhanggutai, China. Forests 10, 954. (doi:10.3390/f10110954). | China | China | 2018 | Restoration | Both | Environmental DNA |
| Halifu, S., Deng, X., Song, X. & Song, R. 2019 Effects of two Trichoderma strains on plant growth, rhizosphere soil nutrients, and fungal community of Pinnus sylvestris var. mongolica annual seedlings. Forests 10, 758. (doi:10.3390/f10090758). | China | China | NR | Restoration | Both | Environmental DNA |
| Han, J.Y., Xiao, H. & Gao, J. 2016 Seasonal dynamics of mycorrhizal fungi in Paphiopedilum spicerianum (Rchb. f) Pfitzer — A critically endangered orchid from China. Global Ecology and Conservation 6, 327-338. (doi:10.1016/j.gecco.2016.03.011). | China | China | 2012 | Conservation | Conservation | Environmental DNA |
| Harrisson, K.A., Magrath, M.J.L., Yen, J.D.L., Pavlova, A., Murray, N., Quin, B., Menkhorst, P., Miller, K.A., Cartwright, K. & Sunnucks, P. 2019 Lifetime fitness costs of inbreeding and being inbred in a critically endangered bird. Current Biology 29, 2711-2717. (doi:10.1016/j.cub.2019.06.064). | Australia | Australia | 1986-2018 | Both | Both | Population genomics |
| Herrera, S. & Shank, T.M. 2016 RAD sequencing enables unprecedented phylogenetic resolution and objective species delimitation in recalcitrant divergent taxa. Molecular Phylogenetics and Evolution 100, 70-79. (doi:10.1016/j.ympev.2016.03.010). | “Global” | USA | 2001-2013 | Conservation | Conservation | Population genomics |
| Hess, J.E., Campbell, N.R., Docker, M.F., Baker, C., Jackson, A., Lampman, R., McIlraith, B., Moser, M.L., Statler, D.P., Young, W.P., et al. 2015 Use of genotyping by sequencing data to develop a high-throughput and multifunctional SNP panel for conservation applications in Pacific lamprey Molecular Ecology Resources 15, 187-202. (doi:10.1111/1755-0998.12283). | “Pacific” | USA | 2007-2012 | Conservation | Conservation | Population genomics |

| | | | | | | |
|---|---|---|---|---|---|---|
| Hohenlohe, P.A., Amish, S.J., Catchen, J.M., Allendorf, F.W. & Luikart, G. 2011 Next-generation RAD sequencing identifies thousands of SNPs for assessing hybridization between rainbow and westslope cutthroat trout Molecular Ecology Resources 11, 117-122. (doi:10.1111/j.1755-0998.2010.02967.x). | USA | USA | NR | Conservation | Conservation | Population genomics |
| Huang, J.-P. 2019 Holocene population decline and conservation implication for the Western Hercules beetle, Dynastes grantii (Coleoptera, Scarabaeidae). Journal of Heredity 110, 629-637. (doi:10.1093/jhered/esz036). | Canada | Taiwan, USA | NR | Conservation | Conservation | Population genomics |
| Huang, Z., Gallot, A., Lao, N.T., Puechmaille, S.J., Foley, N.M., Jebb, D., Bekaert, M. & Teeling, E.C. 2016 A nonlethal sampling method to obtain, generate and assemble whole blood transcriptomes from small, wild mammals. Molecular Ecology Resources 16, 150-162. (doi:10.1111/1755-0998.12447). | France | Ireland, France, Germany | 2013 | Conservation | Conservation | Population genomics |
| Iwanowicz, D.D., Vandergast, A.G., Cornman, R.S., Adams, C.R., Kohn, J.R., Fisher, R.N. & Brehme, C.S. 2016 Metabarcoding of fecal samples to determine herbivore diets: a case study of the endangered Pacific pocket mouse. PLoS One 11, e0165366. (doi:10.1371/journal.pone.0165366). | USA | USA | 2014 | Conservation | Conservation | Environmental DNA |
| Izuddin, M., Srivathsan, A., Lee, A.L., Yam, T.W. & Webb, E.L. 2019 Availability of orchid mycorrhizal fungi on roadside trees in a tropical urban landscape. Scientific Reports 9, 19528. (doi:10.1038/s41598-019-56049-y). | Singapore | Singapore | 2016 | Conservation | Conservation | Environmental DNA |
| Jacobs, A., Carruthers, M., Eckmann, R., Yohannes, E., Adams, C.E., Behrmann-Godel, J. & Elmer, K.R. 2019 Rapid niche expansion by selection on functional genomic variation after ecosystem recovery. Nature Ecology & Evolution 3, 77-86. (doi:10.1038/s41559-018-0742-9). | Germany | UK, Germany | 2014 | Restoration | Both | Population genomics |
| Jahner, J.P., Matocq, M.D., Malaney, J.L., Cox, M., Wolff, P., Gritts, M.A. & Parchman, T.L. 2019 The genetic legacy of 50 years of desert bighorn sheep translocations. Evolutionary Applications 12, 198-213. (doi:10.1111/eva.12708). | USA | USA | 1968-2016 | Conservation | Conservation | Population genomics |
| Janssen, P., Bec, S., Fuhr, M., Taberlet, P., Brun, J.-J. & Bouget, C. 2018 Present conditions may mediate the legacy effect of past land-use changes on species richness and composition of above- and below-ground assemblages. Journal of Ecology 106, 306-318. (doi:10.1111/1365-2745.12808). | France | France | NR | Conservation | Both | Environmental DNA |
| Jia, T., Guo, T., Cao, M. & Chai, B. 2018 Effects of heavy metals on phyllosphere and rhizosphere microbial community of Bothriochloa ischaemum. Applied Science 8, 1419. (doi:10.3390/app8091419). | China | China | 2016 | Restoration | Restoration | Environmental DNA |

| | | | | | | |
|---|---|---|---|---|---|---|
| Jung, J., Philippot, L. & Park, W. 2016 Metagenomic and functional analyses of the consequences of reduction of bacterial diversity on soil functions and bioremediation in diesel-contaminated microcosms. Scientific Reports 6, 23012. (doi:10.1038/srep23012). | Republic of Korea | Republic of Korea | NR | Restoration | Restoration | Environmental DNA |
| Kim, J.-Y., Jeong, S., Kim, K.H., Lim, W.-J., Lee, H.-Y. & Kim, N. 2019 Discovery of genomic characteristics and selection signatures in Korean indigenous goats through comparison of 10 goat breeds. Frontiers in Genetics 10, 699. (doi:10.3389/fgene.2019.00699). | "Europe", "Africa" | South Korea | NR | Conservation | Conservation | Population genomics |
| Kjeldsen, S.R., Raadsma, H.W., Leigh, K.A., Tobey, J.R., Phalen, D., Krockenberger, A., Ellis, W.A., Hynes, E., Higgins, D.P. & Zenger, K.R. 2019 Genomic comparisons reveal biogeographic and anthropogenic impacts in the koala (Phascolarctos cinereus): a dietary-specialist species distributed across heterogeneous environments. Heredity 122, 525-544. (doi:10.1038/s41437-018-0144-4). | Australia | Australia, USA | NR | Conservation | Conservation | Population genomics |
| Kolkert, H., Andrew, R., Smith, R., Rader, R. & Reid, N. 2020 Insectivorous bats selectively source moths and eat mostly pest insects on dryland and irrigated cotton farms. Ecology and Evolution 10, 371-388. (doi:10.1002/ece3.5901). | Australia | Australia | 2012-2014 | Conservation | Conservation | Environmental DNA |
| Kowalczyk, R., Wójcik, J.M., Taberlet, P., Kamiński, T., Miquel, C., Valentini, A., Craine, J.M. & Coissac, E. 2019 Foraging plasticity allows a large herbivore to persist in a sheltering forest habitat: DNA metabarcoding diet analysis of the European bison. Forest Ecology and Management 449, 117474. (doi:10.1016/j.foreco.2019.117474). | Poland | Poland, France, USA | 2008 | Conservation | Conservation | Environmental DNA |
| Kumaresan, D., Cross, A.T., Moreira-Grez, B., Kariman, K., Nevill, P., Stevens, J., Allcock, R.J.N., O'Donnell, A.G., Dixon, K.W. & Whitely, A.S. 2017 Microbial functional capacity is preserved within engineered soil formulations used in mine site restoration. Scientific Reports 7, 564. (doi:10.1038/s41598-017-00650-6). | Australia | Australia | NR | Restoration | Restoration | Environmental DNA |
| Lacoursière-Roussel, A., Dubois, Y., Normandeau, E. & Bernatchez, L. 2016 Improving herpetological surveys in eastern North America using the environmental DNA method. Genome 59, 991-1007. (doi:10.1139/gen-2015-0218). | Canada, USA | Canada | 2013 | Conservation | Conservation | Environmental DNA |
| Lance, A.C., Carrino-Kyker, S.R., Burke, D.J. & Burns, J.H. 2020 Individual plant-soil feedback effects influence tree growth and rhizosphere fungal communities in a temperate forest restoration experiment. Frontiers in Ecology and Evolution 7, 500. (doi:10.3389/fevo.2019.00500). | USA | USA | 2016 | Restoration | Both | Environmental DNA |

| | | | | | | |
|---|---|---|---|---|---|---|
| Lanes, É.C., Pope, N.S., Alves, R., Carvalho Filho, N.M., Giannin, T.C., Giulietti, A.M., Imperatriz-Fonseca, V.L., Monteiro, W., Oliveira, G., Silva, A.R., et al. 2018 Landscape genomic conservation assessment of a narrow-endemic and a widespread morning glory from Amazonian savannas. Frontiers in Plant Science 9, 532. (doi:10.3389/fpls.2018.00532). | Brazil | Brazil, USA, Switzerland | 2016 | Conservation | Both | Population genomics |
| Lavoie, C., Courcelle, M., Redivo, B. & Derome, N. 2018 Structural and compositional mismatch between captive and wild Atlantic salmon (Salmo salar) parrs' gut microbiota highlights the relevance of integrating molecular ecology for management and conservation methods Evolutionary Applications 11, 1671-1685. (doi:10.1111/eva.12658). | Canada | Canada, France, Belgium | 2012-2013 | Conservation | Conservation | Environmental DNA |
| Lecaudey, L.A., Scheletterer, M., Kuzovlev, V.V., Hahn, C. & Weiss, S.J. 2019 Fish diversity assessment in the headwaters of the Volga River using environmental DNA metabarcoding. Aquatic Conservation: Marine and Freshwater Ecosystems 29, 1785-1800. (doi:10.1002/aqc.3163). | Ukraine | Norway, Austria, Russia | 2017 | Conservation | Conservation | Environmental DNA |
| Leitwein, M., Cayuela, H., Ferchaud, A.-L., Normandeau, É., Gagnaire, P.-A. & Bernatchez, L. 2019 The role of recombination on genome-wide patterns of local ancestry exemplified by supplemented brook charr populations. Molecular Ecology 28, 4755-4769. (doi:10.1111/mec.15256). | Canada | Canada, France | 2014-2015 | Conservation | Conservation | Population genomics |
| Leitwein, M., Gagnaire, P.-A., Desmarais, E., Berrebi, P. & Guinand, B. 2018 Genomic consequences of a recent three-way admixture in supplemented wild brown trout populations revealed by local ancestry tracts Molecular Ecology 27, 3466-3483. (doi:10.1111/mec.14816). | France | France | 2015 | Conservation | Conservation | Population genomics |
| Lemay, M.A., Donnelly, D.J. & Russello, M.A. 2013 Transcriptome-wide comparison of sequence variation in divergent ecotypes of kokanee salmon. BMC Genomics 14, 308. (doi:10.1186/1471-2164-14-308). | Canada | Canada | NR | Conservation | Conservation | Population genomics |
| Li, Y., Wu, Z., Dong, X., Wang, D., Qiu, H., Jia, Z. & Sun, Q. 2019 Glucose-induced changes in the bacterial communities of mine tailings at different acidification stages. Canadian Journal of Microbiology 65, 201-213. (doi:10.1139/cjm-2017-0782). | Canada | China | 2015 | Restoration | Restoration | Environmental DNA |
| Li, Y., Chen, L. & Wen, H. 2015 Changes in the composition and diversity of bacterial communities 13 years after soil reclamation of abandoned mine land in eastern China. Ecological Research 30, 357-366. (doi:10.1007/s11284-014-1230-6). | China | China, Malaysia | 2013 | Restoration | Restoration | Environmental DNA |

| | | | | | | |
|---|---|---|---|---|---|---|
| Lim, N.K.M., Tay, Y.C., Srivathsan, A., Tan, J.W.T., Kwik, J.T.B., Baloğlu, B., Meier, R. & Yeo, D.C.J. 2016 Next-generation freshwater bioassessment: eDNA metabarcoding with a conserved metazoan primer reveals species-rich and reservoir-specific communities. Royal Society Open Science 3. (doi:10.1098/rsos.160635). | Singapore | Singapore | NR | Conservation | Conservation | Environmental DNA |
| Limborg, M.T., Larson, W., Shedd, K., Seeb, L.W. & Seeb, J.E. 2017 Novel RAD sequence data reveal a lack of genomic divergence between dietary ecotypes in a landlocked salmonid population. Conservation Genetics Resources 10, 169-171. (doi:10.1007/s12686-017-0791-4 ). | USA | Denmark, USA | 2003-2011 | Conservation | Conservation | Population genomics |
| Liu, G., Chen, L., Shi, Z., Yuan, L.Y., Lock, T.R. & Kallenbach, R.L. 2019 Changes in rhizosphere bacterial and fungal community composition with vegetation restoration in planted forests. Land Degradation & Development 30, 1147-1157. (doi:10.1002/ldr.3275). | China | China, USA | 2015 | Restoration | Restoration | Environmental DNA |
| Liu, Y., Zhang, L., Lu, J., Chen, W., Wei, G. & Lin, Y. 2020 Topography affects the soil conditions and bacterial communities along a restoration gradient on Loess-Plateau. Applied Soil Ecology 150, 103471. (doi:10.1016/j.apsoil.2019.103471). | China | China | 2016 | Restoration | Restoration | Environmental DNA |
| Maaroufi, N.I., Nordin, A., Palmqvist, K., Hasselquist, N.J., Forsmark, B., Rosenstock, N.P., Wallander, H. & Gundale, M.J. 2019 Anthropogenic nitrogen enrichment enhances soil carbon accumulation by impacting saprotrophs rather than ectomycorrhizal fungal activity. Global Change Biology 25, 2900-2914. (doi:10.1111/gcb.14722). | Sweden | Switzerland, Sweden | 2004-2017 | Neither | Both | Environmental DNA |
| Mahon, A.R., Nathan, L.R. & Jerde, C.L. 2014 Meta-genomic surveillance of invasive species in the bait trade. Conservation Genetics Resources 6, 563-567. (doi:10.1007/s12686-014-0213-9). | USA | USA | 2012 | Conservation | Conservation | Environmental DNA |
| Malone, E.W., Perkin, J.S., Leckie, B.M., Kulp, M.A., Hurt, C.R. & Walker, D.M. 2018 Which species, how many, and from where: Integrating habitat suitability, population genomics, and abundance estimates into species reintroduction planning. Global Change Biology 24, 3729-3748. (doi:10.1111/gcb.14126). | USA | USA | NR | Conservation | Conservation | Population genomics |
| Mason, O.U., Scott, N.M., Gonzalez, A., Robbins-Pianka, A., Bælum, J., Kimbrel, J., Bouskill, N.J., Prestat, E., Borglin, S., Joyner, D.C., et al. 2014 Metagenomics reveals sediment microbial community response to Deepwater Horizon oil spill. The ISME Journal 8, 1464-1475. (doi:10.1038/ismej.2013.254). | USA | USA, Denmark, Brazil | 2010 | Conservation | Conservation | Environmental DNA |

| | | | | | | |
|---|---|---|---|---|---|---|
| Massatti, R., Doherty, K.D. & Wood, T.E. 2018 Resolving neutral and deterministic contributions to genomic structure in Syntrichia ruralis (Bryophyta, Pottiaceae) informs propagule sourcing for dryland restoration. Conservation Genetics 19, 85-97. (doi:10.1007/s10592-017-1026-7). | USA | USA | 2012-2013 | Both | Both | Environmental DNA |
| Masse, J., Prescott, C.E., Renaut, S., Terrat, Y. & Grayston, S.J. 2017 Plant community and nitrogen deposition as drivers of alpha and beta diversities of prokaryotes in reconstructed oil sand soils and natural boreal forest soils. Applied and Environmental Microbiology 83, e03319-03316. (doi:10.1128/AEM.03319-16). | Canada | Canada | 2013 | Restoration | Restoration | Environmental DNA |
| McCartney-Melstad, E., Gidiş, M. & Shaffer, H.B. 2018 Population genomic data reveal extreme geographic subdivision and novel conservation actions for the declining foothill yellow-legged frog. Heredity 121, 112-125. (doi:10.1038/s41437-018-0097-7). | USA | USA, Turkey | NR | Conservation | Conservation | Population genomics |
| McCartney-Melstad, E., Vu, J.K. & Shaffer, H.B. 2018 Genomic data recover previously undetectable fragmentation effects in an endangered amphibian. Molecular Ecology 27, 4430-4443. (doi:10.1111/mec.14892). | USA | USA | 2013-2015 | Conservation | Conservation | Population genomics |
| McInerney, P.J., Shackleton, M.E., Rees, G.N., Frechette, J.L., Sam, H. & Hor, L. 2019 Release of critically endangered crocodiles: Development and application of a food web approach to determine suitability of release habitat. Aquatic Conservation: Marine and Freshwater Ecosystems 29, 1849-1862. (doi:10.1002/aqc.3159). | Cambodia | Australia, Cambodia | 2017 | Conservation | Conservation | Environmental DNA |
| Mikryukov, V.S., Dulya, O.V. & Modorov, M.V. 2020 Phylogenetic signature of fungal response to long-term chemical pollution. Soil Biology and Biochemistry 140, 107644. (doi:10.1016/j.soilbio.2019.107644). | Russia | Russia | 2014 | Restoration | Restoration | Environmental DNA |
| Ming, Y., Jian, J., Yu, X., Wang, J. & Liu, W. 2019 The genome resources for conservation of Indo-Pacific humpback dolphin, Sousa chinensis. Scientific Data 6, 68. (doi:10.1038/s41597-019-0078-6). | China | China | 2015 | Conservation | Conservation | Comparative genomics |
| Moeller, A.H., Peeters, M., Ayouba, A., Ngole, E.M., Esteban, A., Hahn, B.H. & Ochman, H. 2015 Stability of the gorilla microbiome despite simian immunodeficiency virus infection. Molecular Ecology 24, 690-697. (doi:10.1111/mec.13057). | Cameroon | USA, France, Cameroon | 2004-2012 | Conservation | Conservation | Environmental DNA |
| Muniz, F.L., Campos, Z., Hernández Rangel, S.M., Martínez, J.G., Souza, B.C., De Thoisy, B., Botero-Arias, R., Hrbek, T. & Farias, I.P. 2018 Delimitation of evolutionary units in Cuvier's dwarf caiman, Paleosuchus palpebrosus (Cuvier, 1807): insights from conservation | South America | Brazil, Colombia, French Guiana, USA | NR | Conservation | Conservation | Population genomics |

| | | | | | | |
|---|---|---|---|---|---|---|
| of a broadly distributed species. Conservation Genetics 19, 599-610. (doi:10.1007/s10592-017-1035-6). | | | | | | |
| Nathan, L.R., Jerde, C.L., Budny, M.L. & Mahon, A.R. 2015 The use of environmental DNA in invasive species surveillance of the Great Lakes commercial bait trade. Conservation Biology 29, 430-439. (doi:10.1111/cobi.12381). | USA, Canada | USA | 2012 | Conservation | Conservation | Environmental DNA |
| Ning, C., Xiang, W., Mueller, G.M., Egerton-Warburton, L.M., Yan, W. & Liu, S. 2020 Differences in ectomycorrhizal community assembly between native and exotic pines are reflected in their enzymatic functional capacities. Plant and Soil 446, 179-193. (doi:10.1007/s11104-019-04355-9). | China | China, USA | 2016 | Restoration | Restoration | Environmental DNA |
| Nunez, J.C.B., Seale, T.P., Fraser, M.A., Burton, T.L., Fortson, T.N., Hoover, D., Travis, J., Oleksiak, M.F. & Crawford, D.L. 2015 Population genomics of the euryhaline teleost Poecilia latipinna PLoS One 10, E0137077. (doi:10.1371/journal.pone.0137077). | USA | USA | 2014 | Conservation | Conservation | Population genomics |
| Oguntuase, B.G., Ogunjemite, B.G. & Meisel, R.P. 2018 Morphometric and genetic differentiation among populations of flat-headed cusimanse (Crossarchus platycephalus) in Nigeria. Ecology and Evolution 8, 7228-7235. (doi:10.1002/ece3.4262). | Nigeria | Nigeria, USA | NR | Conservation | Conservation | Population genomics |
| Pansu, J., Guyton, J.A., Potter, A.B., Atkins, J.L., Daskin, J.H., Wursten, B., Kartzinel, T.R. & Pringle, R.M. 2019 Trophic ecology of large herbivores in a reassembling African ecosystem. Journal of Ecology 107, 1355-1376. (doi:10.1111/1365-2745.13113). | Mozambique | USA, Belgium | 2016 | Conservation | Conservation | Environmental DNA |
| Pedrinho, A., Mendes, L.W., Merloti, L.F., da Fonseca, M.C., Cannavan, F.S. & Tsai, S.M. 2019 Forest-to-pasture conversion and recovery based on assessment of microbial communities in Eastern Amazon rainforest. FEMS Microbiology Ecology 95. (doi:10.1093/femsec/fiy236). | Brazil | Brazil | 2016 | Restoration | Restoration | Environmental DNA |
| Perry, G.H., Louis Jr, E.E., Ratan, A., Bedoya-Reina, O.C., Burhans, R.C., Lei, R., Johnson, S.E., Schuster, S.C. & Miller, W. 2013 Aye-aye population genomic analyses highlight an important center of endemism in northern Madagascar. Proceedings of the National Academy of Science 110, 5823-5828. (doi:10.1073/pnas.1211990110). | Madagascar | USA, Canada, Singapore | NR | Conservation | Conservation | Population genomics |
| Pilloni, G., Bayer, A., Ruth-Anneser, B., Fillinger, L., Engle, M., Griebler, C. & Lueders, T. 2019 Dynamics of hydrology and anaerobic hydrocarbon degrader communities in a tar-oil contaminated aquifer. Microorganisms 7, 46. (doi:10.3390/microorganisms7020046). | Germany | Germany | 2006 | Restoration | Restoration | Environmental DNA |

| | | | | | | |
|---|---|---|---|---|---|---|
| Pinho, C.J., Santos, B., Mata, V.A., Seguro, M., Romeiras, M.M., Lopes, R.J. & Vasconcelos, R. 2018 What is the giant wall gecko having for dinner? Conservation genetics for guiding reserve management in Cabo Verde. Genes (Basel) 9, 599. (doi:10.3390/genes9120599). | Republic of Cabo Verde | Portugal | 2016-2017 | Conservation | Conservation | Environmental DNA |
| Poorten, T.J. & Rosenblum, E.B. 2016 Comparative study of host response to chytridiomycosis in a susceptible and a resistant toad species. Molecular Ecology 25, 5663-5679. (doi:10.1111/mec.13871). | USA | USA | NR | Conservation | Conservation | Population genomics |
| Porazinska, D.L., Fujisaki, I., Purcell, M.F. & Giblin-Davis, R.M. 2014 Plant invasions from a belowground nematocentric perspective. Soil Biology and Biochemistry 77, 213-220. (doi:10.1016/j.soilbio.2014.06.004). | USA, Australia | USA, Australia | 2008-2010 | Restoration | Restoration | Environmental DNA |
| Port, J.A., O'Donnell, J.L., Romero-Maraccini, O.C., Leary, P.R., Litvin, S.Y., Nickols, K.J., Yamahara, K.M. & Kelly, R.P. 2016 Assessing vertebrate biodiversity in a kelp forest ecosystem using environmental DNA. Molecular Ecology 25, 527-541. (doi:10.1111/mec.13481). | USA | USA | 2013 | Conservation | Conservation | Environmental DNA |
| Purahong, W., Wubet, T., Krüger, D. & Buscot, F. 2019 Application of next-generation sequencing technologies to conservation of wood-inhabiting fungi Conservation Biology 33, 716-724. (doi:10.1111/cobi.13240). | Germany | Germany | 2012 | Conservation | Conservation | Environmental DNA |
| Razgour, O., Forester, B., Taggart, J.B., Bekaert, M., Juste, J., Ibáñez, C., Puechmaille, S.J., Novella-Fernandez, R., Alberdi, A. & Manel, S. 2019 Considering adaptive genetic variation in climate change vulnerability assessment reduces species range loss projections. Proceedings of the National Academy of Science 116, 10418-10423. (doi:10.1073/pnas.1820663116). | Spain | UK, USA, Spain, France, Germany, Denmark | 2010-2015 | Conservation | Conservation | Population genomics |
| Reitzel, A.M., Herrera, S., Layden, M.J., Martindale, M.Q. & Shank, T.M. 2013 Going where traditional markers have not gone before: utility of and promise for RAD sequencing in marine invertebrate phylogeography and population genomics Molecular Ecology 22, 2953-2970. (doi:10.1111/mec.12228). | USA | USA | 2003-2004 | Conservation | Conservation | Population genomics |
| Rosales, S.M., Miller, M.W., Williams, D.E., Traylor-Knowles, N., Young, B. & Serrano, X.M. 2019 Microbiome differences in disease-resistant vs. susceptible Acropora corals subjected to disease challenge assays. Scientific Reports 9, 18279. (doi:10.1038/s41598-019-54855-y). | USA | USA | 2017 | Conservation | Conservation | Environmental DNA |
| Rossetto, M., Bragg, J., Kilian, A., McPherson, H., van der Merwe, M. & Wilson, P.D. 2019 Restore and Renew: a genomics-era framework for species provenance delimitation. Restoration Ecology 27, 538-548. (doi:10.1111/rec.12898). | Australia | Australia | NR | Restoration | Restoration | Population genomics |

| | | | | | | |
|---|---|---|---|---|---|---|
| Schmelzle, M.C. & Kinziger, A.P. 2016 Using occupancy modelling to compare environmental DNA to traditional field methods for regional-scale monitoring of an endangered aquatic species. Molecular Ecology Resources 16, 895-908. (doi:10.1111/1755-0998.12501). | USA | USA | 2014 | Conservation | Both | Environmental DNA |
| Schmidt, D.A., Campbell, N.R., Govindarajulu, P., Larsen, K.W. & Russello, M.A. 2020 Genotyping-in-Thousands by sequencing (GT-seq) panel development and application to minimally invasive DNA samples to support studies in molecular ecology. Molecular Ecology Resources 20, 114-124. (doi:10.1111/1755-0998.13090). | Canada, USA | Canada | NR | Conservation | Conservation | Population genomics |
| Schmidt, D.A., Fallon, S., Roberts, D.T., Espinoza, T., McDougall, A., Brooks, S.G., Kind, P.K., Bond, N.R., Kennard, M.J. & Hughes, J.M. 2018 Monitoring age-related trends in genomic diversity of Australian lungfish. Molecular Ecology 27, 3231-3241. (doi:10.1111/mec.14791). | Australia | Australia | 2013-2015 | Conservation | Conservation | Population genomics |
| Seeber, P.A., McEwen, G.K., Löber, U., Förster, D.W., East, M.L., Melzheimer, J. & Greenwood, A.D. 2019 Terrestrial mammal surveillance using hybridization capture of environmental DNA from African waterholes. Molecular Ecology Resources 19, 1486-1496. (doi:10.1111/1755-0998.13069). | Namibia, Tanzania | Germany | 2015-2016 | Conservation | Conservation | Environmental DNA |
| Siddall, M.E., Barkdull, M., Tessler, M., Brugler, M.R., Borda, E. & Hekkala, E. 2019 Ideating iDNA: Lessons and limitations from leeches in legacy collections. PLoS One 14, e0212226. (doi:10.1371/journal.pone.0212226). | China | USA | 2014-2015 | Conservation | Conservation | Environmental DNA |
| Sigsgaard, E.E., Torquato, F., Frøslev, T.G., Moore, A.B.M., Sørensen, J.M., Range, P., Ben-Hamadou, R., Bach, S.S., Møller, P.R. & Thomsen, P.F. 2019 Using vertebrate environmental DNA from seawater in biomonitoring of marine habitats. Conservation Biology 34, 697-710. (doi:10.1111/cobi.13437 ). | Arabian Gulf | Denmark, UK, Qatar | 2016-2017 | Conservation | Both | Environmental DNA |
| Singh, A., Lasek-Nesselquist, E., Chaturvedi, V. & Chaturvedi, S. 2018 Trichoderma polysporum selectively inhibits white-nose syndrome fungal pathogen Pseudogymnoascus destructans amidst soil microbes. Microbiome 6, 139. (doi:10.1186/s40168-018-0512-6). | USA | USA | 2015 | Conservation | Conservation | Environmental DNA |
| Smart, M.D., Cornman, R.S., Iwanowicz, D.D., McDermott-Kubeczko, M., Pettis, J.S., Spivak, M.S. & Otto, C.R.V. 2017 A comparison of honey bee-collected pollen from working agricultural lands using light microscopy and ITS metabarcoding. Environmental Entomology 46, 38-49. (doi:10.1093/ee/nvw159). | USA | USA | 2010-2011 | Conservation | Both | Environmental DNA |

| | | | | | | |
|---|---|---|---|---|---|---|
| Srivathsan, A., Ang, A., Vogler, A.P. & Meier, R. 2016 Fecal metagenomics for the simultaneous assessment of diet, parasites, and population genetics of an understudied primate. Frontiers in Zoology 13, 17. (doi:10.1186/s12983-016-0150-4). | Singapore | Singapore, USA, UK | 2008-2011 | Conservation | Conservation | Environmental DNA |
| Steane, D.A., Mclean, E.H., Potts, B.M., Prober, S.M., Stock, W.D., Stylianou, V.M., Vaillancourt, R.E. & Byrne, M. 2017 Evidence for adaptation and acclimation in a widespread eucalypt of semi-arid Australia Biological Journal of the Linnean Society 121, 484-500. (doi:10.1093/biolinnean/blw051). | Australia | Australia | 2011 | Restoration | Both | Population genomics |
| Sun, S. & Badgley, B.D. 2019 Changes in microbial functional genes within the soil metagenome during forest ecosystem restoration. Soil Biology and Biochemistry 135, 163-172. (doi:10.1016/j.soilbio.2019.05.004). | USA | USA | 2014 | Restoration | Restoration | Environmental DNA |
| Sun, S., Li, S., Avera, B.N., Strahm, B.D. & Badgley, B.D. 2017 Soil bacterial and fungal communities show distinct recovery patterns during forest ecosystem restoration. Applied and Environmental Microbiology 83, e00966-00917. (doi:10.1128/AEM.00966-17). | USA | USA | 2013 | Restoration | Restoration | Environmental DNA |
| Sun, Y., Liu, Z., Zhang, Y., Lai, Z., She, W., Bai, Y., Feng, W. & Qin, S. 2020 Microbial communities and their genetic repertoire mediate the decomposition of soil organic carbon pools in revegetation shrublands in a desert in northern China. European Journal of Soil Science 71, 93-105. (doi:10.1111/ejss.12824). | China | China | 2016 | Restoration | Restoration | Environmental DNA |
| Taylor, R.S., Bolton, M., Beard, A., Birt, T., Deane-Coe, P., Raine, A.F., González-Solís, J., Lougheed, S.C. & Friesen, V.L. 2019 Cryptic species and independent origins of allochronic populations within a seabird species complex (Hydrobates spp.). Molecular Phylogenetics and Evolution 139, 106552. (doi:10.1016/j.ympev.2019.106552). | “Oceans” | Canada, UK, Ascension and Tristan de Cunha, Spain | NR | Conservation | Conservation | Population genomics |
| Trivedi, J., Lachapelle, J., Vanderwolf, K.J., Misra, V., Willis, C.K.R., Ratcliffe, J.M., Ness, R.W., Anderson, J.B. & Kohn, L.M. 2017 Fungus causing white-nose syndrome in bats accumulates genetic variability in North America with no sign of recombination. mSphere 2, e00271-00217. (doi:10.1128/mSphereDirect.00271-17). | USA | Canada | NR | Conservation | Conservation | Comparative genomics |
| Thomsen, P.F., Kielgast, J., Iversen, L.L., Wiuf, C., Rasmussen, M., Gilbert, M.T.P., Orlando, L. & Willerslev, E. 2012 Monitoring endangered freshwater biodiversity using environmental DNA. Molecular Ecology 21, 2565-2573. (doi:10.1111/j.1365-294X.2011.05418.x). | “Europe” | Denmark | 2009 | Conservation | Both | Environmental DNA |

| | | | | | | |
|---|---|---|---|---|---|---|
| Tilker, A., Abrams, J.F., Nguyen, A., Hörig, L., Axtner, J., Louvrier, J., Rawson, B.M., Nguyen, H.A.Q., Guegan, F., Nguyen, T.V., et al. 2020 Identifying conservation priorities in a defaunated tropical biodiversity hotspot. Diversity and Distributions 26, 426-440. (doi:10.1111/ddi.13029). | Vietnam, Laos | Germany, USA, Vietnam, Lao PDR | 2011-2017 | Conservation | Both | Environmental DNA |
| Valentini, A., Taberlet, P., Miaud, C., Civade, R., Herder, J., Thomsen, P.F., Bellemain, E., Besnard, A., Coissac, E., Boyer, F., et al. 2016 Next-generational monitoring of aquatic biodiversity using environmental DNA metabarcoding. Molecular Ecology 25, 929-942. (doi:10.1111/mec.13428). | “Europe” | France, Netherlands, Denmark, UK, Canada | 2011-2014 | Conservation | Both | Environmental DNA |
| van der Valk, T., Gonda, C.M., Silegowa, H., Almanza, S., Sifuentes-Romero, I., Hart, T.B., Hart, J.A., Detwilder, K.M. & Guschanski, K. 2020 The genome of the endangered dryas monkey provides new insights into the evolutionary history of the vervets. Molecular Biology and Evolution 37, 183-194. (doi:10.1093/molbev/msz213). | Democratic Republic of Congo | Sweden, Democratic Republic of Congo, USA | 2014-2017 | Conservation | Conservation | Comparative genomics |
| Vega, R., Vázquez-Domínguez, E., White, T.A., Valenzuela-Galván, D. & Searle, J.B. 2017 Population genomics applications for conservation: the case of the tropical dry forest dweller Peromyscus melanophrys. Conservation Genetics 18, 313-326. (doi:10.1007/s10592-016-0907-5). | USA | UK, USA, Mexico | 2002-2011 | Conservation | Conservation | Population genomics |
| Vermeulen, E.T., Lott, M.J., Eldridge, M.D.B. & Power, M.L. 2016 Parasites on the hop: Captive breeding maintains biodiversity of Eimeria communities in an endangered marsupial. Biological Conservation 200, 17-25. (doi:10.1016/j.biocon.2016.05.019). | Australia | Australia | 2009-2014 | Conservation | Conservation | Environmental DNA |
| Viana, J.P.G., Siqueira, M.V.B.M., Araujo, F.L., Grando, C., Sujii, P.S., Silvestre, E.d.A., Novello, M., Pinheiro, J.B., Cavallari, M.M., Brancalion, P.H.S., et al. 2018 Genomic diversity is similar between Atlantic Forest restorations and natural remnants for the native tree Casearia sylvestris Sw. PLoS One 13, e0192165. (doi:10.1371/journal.pone.0192165). | Brazil | Brazil, USA | NR | Both | Both | Population genomics |
| von Thaden, A., Nowak, C., Tiesmeyer, A., Reiners, T.E., Alves, P.C., Lyons, L.A., Mattucci, F., Randi, E., Cragnolini, M., Galián, J., et al. 2020 Applying genomic data in wildlife monitoring: Development guidelines for genotyping degraded samples with reduced single nucleotide polymorphism panels Molecular Ecology Resources 20, 662-680. (doi:10.1111/1755-0998.13136). | Germany | Germany, Portugal, USA, Italy, Denmark, Spain, Romania, UK, Belgium | 2001-2016 | Conservation | Conservation | Population genomics |
| Wang, C., Zhang, W., Zhao, C., Shi, R., Xue, R. & Li, X. 2020 Revegetation by sowing reduces soil bacterial and fungal diversity. Ecology and Evolution 10, 431-440. (doi:10.1002/ece3.5906). | China | China | 2017-2018 | Restoration | Restoration | Environmental DNA |

| | | | | | | |
|---|---|---|---|---|---|---|
| Wang, X., Hua, F., Wang, L., Wilcove, D.S. & Yu, D.W. 2019 The biodiversity benefit of native forests and mixed-species plantations over monoculture plantations. Diversity and Distributions 25, 1721-1735. (doi:10.1111/ddi.12972). | China | China, USA, UK | NR | Both | Both | Environmental DNA |
| Wang, Z., Chen, Q., Liu, L., Wen, X. & Liao, Y. 2016 Responses of soil fungi to 5-year conservation tillage treatments in the drylands of northern China. Applied Soil Ecology 101, 132-140. (doi:10.1016/j.apsoil.2016.02.002). | China | China | 2014 | Conservation | Both | Environmental DNA |
| Waterhouse, B.R., Boyer, S. & Wratten, S.D. 2014 Pyrosequencing of prey DNA in faeces of carnivorous land snails to facilitate ecological restoration and relocation programmes. Oecologia 175, 737-746. (doi:10.1007/s00442-014-2933-7). | New Zealand | New Zealand | 2011-2012 | Both | Both | Environmental DNA |
| Wei, Z., Hao, Z., Li, X., Guan, Z., Cai, Y. & Liao, X. 2019 The effects of phytoremediation on soil bacterial communities in an abandoned mine site of rare earth elements. Science of the Total Environment 670, 950-960. (doi:10.1016/j.scitotenv.2019.03.118). | China | China | 2016 | Restoration | Restoration | Environmental DNA |
| Wicaksono, C.Y., Aguirre-Guiterrez, J., Nouhra, E., Pastor, N., Raes, N., Pacheco, S. & Geml, J. 2017 Contracting montane cloud forests: a case study of the Andean alder (Alnus acuminata) and associated fungi in the Yungas Biotropica 49, 141-152. (doi:10.1111/btp.12394). | Argentina | Netherlands, Argentina | NR | Conservation | Both | Environmental DNA |
| Wu, W., Ng, W.-L., Yang, J.-X., Li, W.-M. & Ge, X.-J. 2018 High cryptic species diversity is revealed by genome-wide polymorphisms in a wild relative of banana, Musa itinerans, and implications for its conservation in subtropical China. BMC Plant Biology 18, 194. (doi:10.1186/s12870-018-1410-6). | China | China | NR | Conservation | Conservation | Population genomics |
| Xiong, M., Wang, D., Bu, H., Shao, X., Zhang, D., Li, S., Wang, R. & Yao, M. 2017 Molecular dietary analysis of two sympatric felids in the Mountains of Southwest China biodiversity hotspot and conservation implications. Scientific Reports 7, 41909. (doi:10.1038/srep41909). | China | China | 2013-2014 | Conservation | Conservation | Environmental DNA |
| Xiong, W., Ni, P., Chen, Y., Gao, Y., Li, S. & Zhan, A. 2019 Biological consequences of environmental pollution in running water ecosystems: A case study in zooplankton. Environmental Pollution 252, 1483-1490. (doi:10.1016/j.envpol.2019.06.055). | China | China | 2017 | Both | Both | Environmental DNA |
| Xu, B., Sun, G., Wang, X., Lu, J., Wang, I.J. & Wang, Z. 2017 Population genetic structure is shaped by historical, geographic, and environmental factors in the leguminous shrub Caragana microphylla on the Inner Mongolia Plateau of China. BMC Plant Biology 17, 200. (doi:10.1186/s12870-017-1147-7). | China | USA, China | NR | Conservation | Both | Population genomics |

| | | | | | | |
|---|---|---|---|---|---|---|
| Yan, D., Bissett, A., Gellie, N., Mills, J.G., Lowe, A.J. & Breed, M.F. 2020 Soil bacterial community differences along a coastal restoration chronosequence. Plant Ecology 221, 795-811. (doi:10.1007/s11258-019-00979-0). | Australia | China, Australia | 2015 | Restoration | Restoration | Environmental DNA |
| Yan, D., Gellie, N.J.C., Mills, J.G., Connell, G., Bissett, A., Lowe, A.J. & Breed, M.F. 2020 A soil archael community responds to a decade of ecological restoration. Restoration Ecology 28, 63-72. (doi:10.1111/rec.13033). | Australia | China, Australia | 2015 | Restoration | Restoration | Environmental DNA |
| Yan, D., Mills, J.G., Gellie, N.J.C., Bissett, A., Lowe, A.J. & Breed, M.F. 2018 High-throughput eDNA monitoring of fungi to track functional recovery in ecological restoration. Biological Conservation 217, 113-120. (doi:10.1016/j.biocon.2017.10.035). | Australia | China, Australia | 2015 | Restoration | Restoration | Environmental DNA |
| Yang, C., Wang, X., Miller, J.A., de Blécourt, M., Ji, Y., Yang, C., Harrison, R.D. & Yu, D.W. 2014 Using metabarcoding to ask if easily collected soil and leaf-litter samples can be used as a general biodiversity indicator. Ecological Indicators 46, 379-389. (doi:10.1016/j.ecolind.2014.06.028). | China | China, Netherlands, Germany, UK | 2010-2012 | Both | Both | Environmental DNA |
| Yang, Y., Cheng, H., Liu, L., Dou, Y. & An, S. 2020 Comparison of soil microbial community between planted woodland and natural grass vegetation on the Loess Plateau. Forest Ecology and Management 460, 117817. (doi:10.1016/j.foreco.2019.117817). | China | China, USA | 2017 | Restoration | Restoration | Environmental DNA |
| Yang, Y., Gao, Y., Chen, Y., Li, S. & Zhan, A. 2019 Interactome-based abiotic and biotic impacts on biodiversity of plankton communities in disturbed wetlands. Diversity and Distributions 25, 1416-1428. (doi:10.1111/ddi.12949). | China | China | NR | Both | Both | Environmental DNA |
| Yao, R., Xu, L., Hu, T., Chen, H., Qi, D., Gu, X., Yang, X., Yang, Z. & Zhu, L. 2019 The “wildness” of the giant panda gut microbiome and its relevance to effective translocation. Global Ecology and Conservation 18, e00644. (doi:10.1016/j.gecco.2019.e00644). | China | China | 2012-2014 | Conservation | Conservation | Environmental DNA |
| Yates, M.C., Bowles, E. & Fraser, D.J. 2019 Small population size and low genomic diversity have no effect on fitness in experimental translocations of a wild fish. Proceedings of the Royal Society B 286, 20191989. (doi:10.1098/rspb.2019.1989). | Canada | China | 201202015 | Conservation | Conservation | Population genomics |
| Zadodna, M., Grueber, C.E. & Gemmell, N.J. 2013 Parallel tagged next-generation sequencing on pooled samples – A new approach for population genetics in ecology and conservation. PLoS One 8, e61471. (doi:10.1371/journal.pone.0061471). | New Zealand | New Zealand | NR | Conservation | Both | Environmental DNA |

| | | | | | | |
|---|---|---|---|---|---|---|
| Zampolli, J., Di Canito, A., Cappelletti, M., Collina, E., Lasagni, M. & Di Gennaro, P. 2020 Biodegradation of naphthenic acids: identification of Rhodococcus opacus R7 genes as molecular markers for environmental monitoring and their application in slurry microcosms. Applied Microbiology and Biotechnology 104, 2675-2689. (doi:10.1007/s00253-020-10378-5). | Italy | Italy | NR | Restoration | Restoration | Environmental DNA |
| Zimmerman, S., Aldridge, C.L., Oh, K.P., Cornman, R.S. & Oyler-McCance, S.J. 2019 Signatures of adaptive divergence among populations of an avian species of conservation concern. Evolutionary Applications 12, 1661-1677. (doi:10.1111/eva.12825). | USA | USA | 1996-2004 | Conservation | Conservation | Population genomics |
| Zou, K., Chen, J., Ruan, H., Li, Z., Guo, W., Li, M. & Liu, L. 2020 eDNA metabarcoding as a promising conservation tool for monitoring fish diversity in a coastal wetland of the Pearl River Estuary compared to bottom trawling. Science of the Total Environment 702, 134704. (doi:10.1016/j.scitotenv.2019.134704). | China | China | 2017-2018 | Conservation | Both | Environmental DNA |

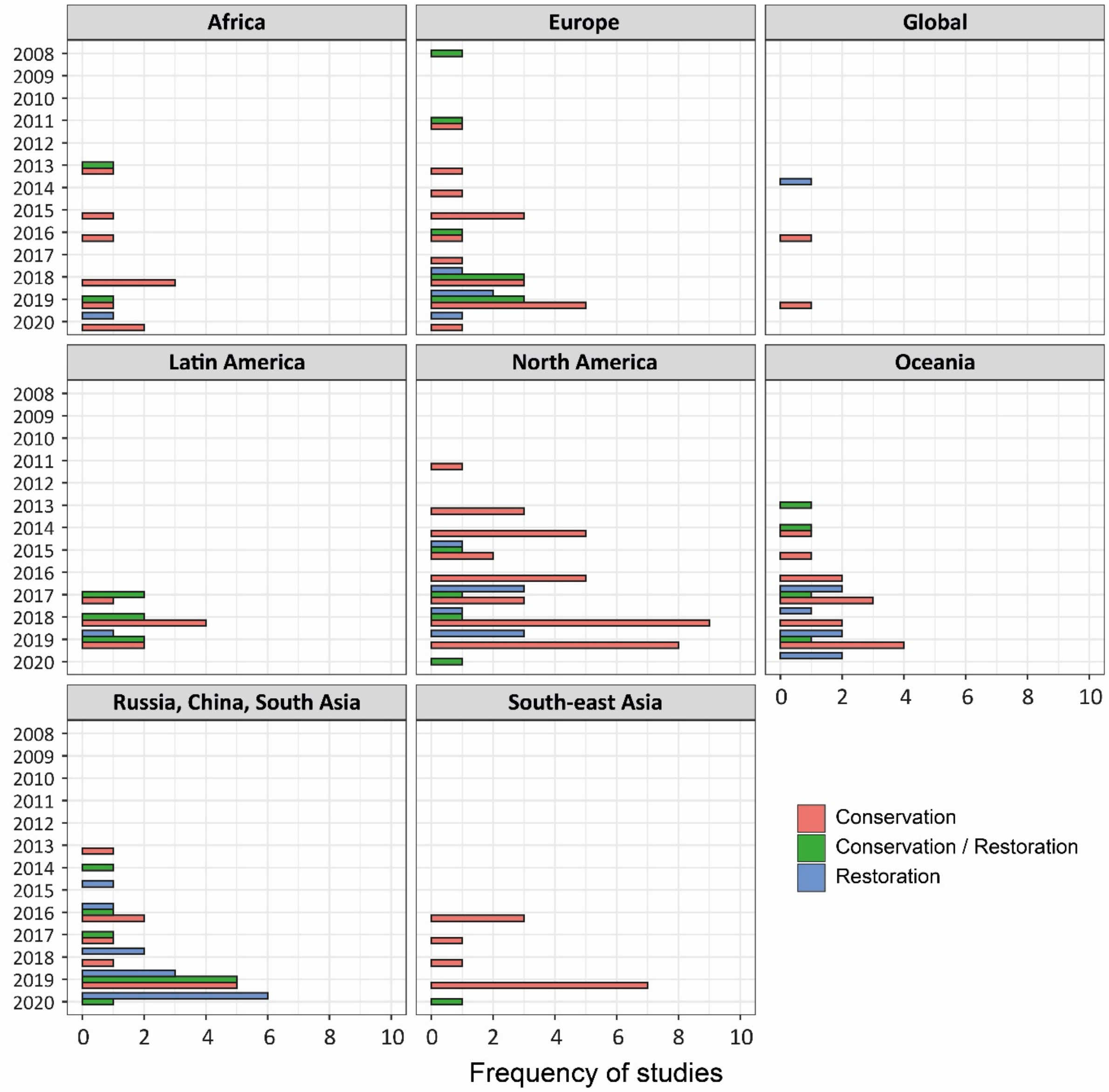


**Appendix 5.** Frequency of published studies from the systematic mapping review by geographic region, across time. For each region, studies were divided into conservation (orange), conservation/restoration (green), and restoration (blue).
